# Flexible linkers in CaMKII control the balance between activating and inhibitory autophosphorylation

**DOI:** 10.1101/848044

**Authors:** Moitrayee Bhattacharyya, Young Kwang Lee, Serena Muratcioglu, Baiyu Qiu, Priya Nyayapati, Howard Schulman, Jay Groves, John Kuriyan

## Abstract

The activity of Ca^2+^/calmodulin-dependent protein kinase II (CaMKII) depends on the balance between activating and inhibitory autophosphorylation (Thr 286 and Thr 305/306, respectively, in the human α isoform). Variation in the lengths of the flexible linkers that connect the kinase domains of CaMKII to a central oligomeric hub could alter transphosphorylation rates within a holoenzyme, thereby affecting the balance of autophosphorylation outcomes. Using a single-molecule assay for visualization of CaMKII phosphorylation on glass, we show that the balance of autophosphorylation is flipped between CaMKII-α and CaMKII-β, the two principal isoforms in the brain. CaMKII-α, with a ∼30 residue kinase-hub linker, readily acquires activating autophosphorylation, which we show is resistant to removal by phosphatases. CaMKII-β, with a ∼200 residue kinase-hub linker, is biased towards inhibitory autophosphorylation. Thus, the responsiveness of CaMKII to calcium signals can be tuned by varying the relative levels of the α and β isoforms.

## Introduction

A characteristic feature of signaling by protein kinases is that the kinases are themselves regulated by phosphorylation. For most kinases, this involves the phosphorylation of one or more residues in the activation loop, a regulatory element located at the active site of the kinase (Huse and Kuriyan, 2002; Johnson et al., 1996; Nolen et al., 2004). An exception is provided by Ca^2+^/calmodulin-dependent protein kinase II (CaMKII), an enzyme that has crucial roles in animal-cell signaling, particularly in the brain and in cardiac tissue (Bayer and Schulman, 2019; Bhattacharyya et al., 2019; Griffith, 2004; Hudmon and Schulman, 2002; Kennedy, 2013; Lisman et al., 2012; Stratton et al., 2013). CaMKII has no phosphorylation sites within its activation loop. Instead, the activity of CaMKII is modulated by autophosphorylation on two sites, Thr 286 and Thr 305/306 (residue numbering corresponds to the sequence of human CaMKII-α), both located in a regulatory segment that immediately follows the kinase domain (Figure 1a, d).

**Figure 1:**
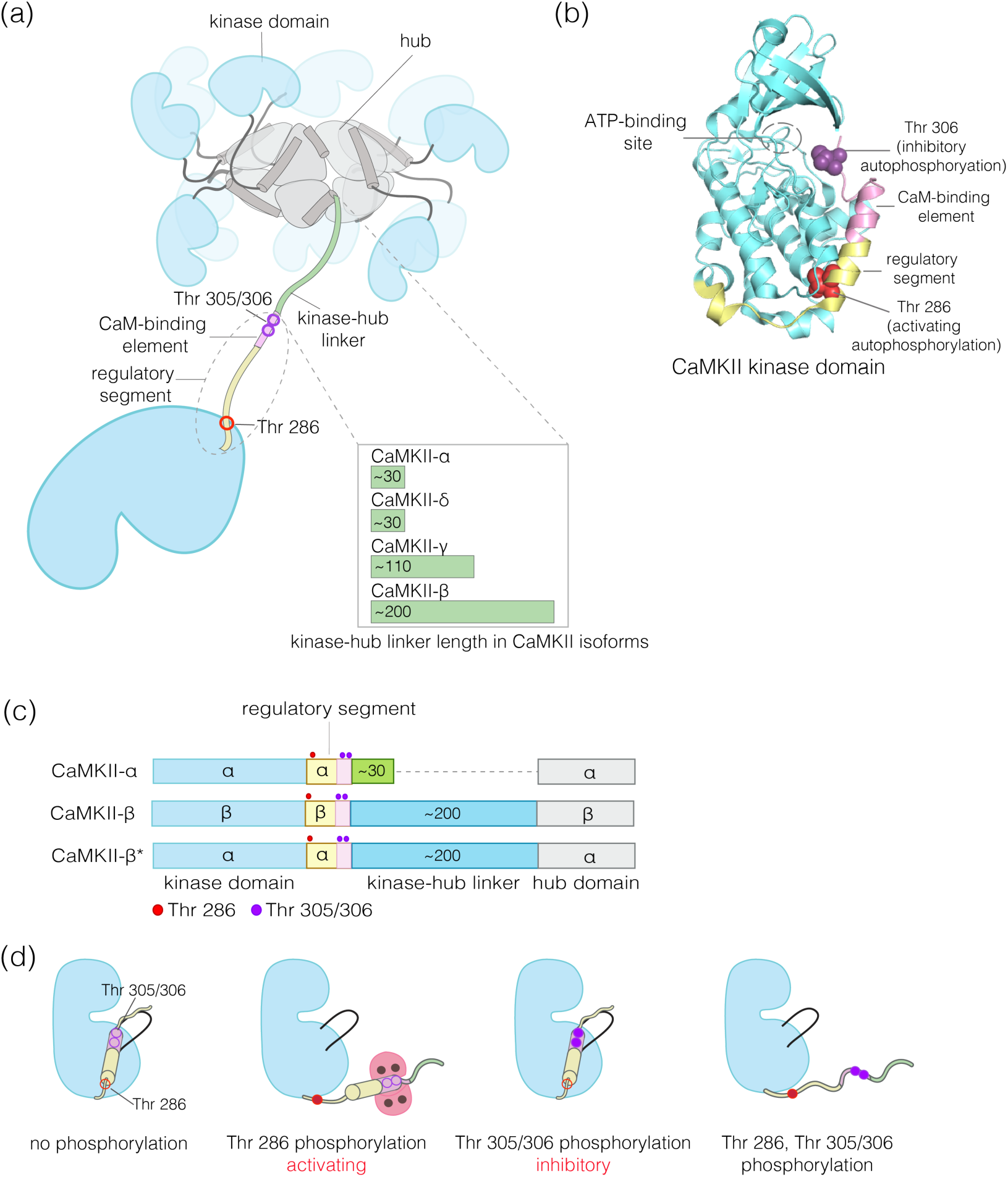
Structural organization and Ca^2+^/CaM-dependent activation of CaMKII. (a) CaMKII is organized as a holoenzyme with kinase domains connected to a central dodecameric/tetradecameric hub by a regulatory segment and a flexible linker, referred to as the kinase-hub linker. All the domains are labeled and this color scheme used will be maintained throughout. The kinase-hub linker is the principle difference between the four CaMKII isoforms: α/β/γ/δ. (b) Crystal structure of the autoinhibited kinase domain from human CaMKII-δ with a small molecule inhibitor (PDB ID: 2VN9) bound (Rellos et al. 2010). The regulatory segment places Thr 306 optimally for *cis*-phosphorylation, while Thr 286 at the base of the kinase can only be phosphorylated in *trans*. (c) Schematic diagram showing the design principle for CaMKII-β*, a surrogate for CaMKII-β. (d) Depiction of the four possible phosphorylation states of CaMKII.

Phosphorylation of Thr 286 confers Ca^2+^/calmodulin-independent activity, referred to as autonomy (Lai et al., 1986; Lou et al., 1986; Miller et al., 1988; Miller and Kennedy, 1986; Saitoh and Schwartz, 1985; Schworer et al., 1986). We refer to Thr 286 phosphorylation as activating phosphorylation, since it prolongs the active state. In contrast, phosphorylation on Thr 305/306 is inhibitory, because it blocks the binding of Ca^2+^/CaM (Colbran, 1993; Hanson and Schulman, 1992; Hashimoto et al., 1987; Lou and Schulman, 1989; Patton et al., 1990). Mutation of these autophosphorylation sites results in profound alterations in learning and memory (Elgersma et al., 2002; Giese et al., 1998; Giese and Mizuno, 2013; Küry et al., 2017; Silva et al., 1992).

CaMKII is organized into a dodecameric or tetradecameric holoenzyme consisting of a central ring-shaped hub assembly, to which the kinase domains are connected by flexible linkers (the kinase-hub linkers) (Figure 1a) (Chao et al., 2011; Myers et al., 2017; Rosenberg et al., 2005). The four isoforms of human CaMKII, denoted α, β, γ, and δ, and their ∼40 splice variants, differ principally in the length and sequence of the kinase-hub linkers. The kinase and the hub domains of these isoforms share about 90% and 80% sequence identity, respectively (Tombes et al., 2003). CaMKII-α and CaMKII-δ have linkers spanning 31 residues each, while CaMKII-β and CaMKII-γ have longer linkers (218 and 110 residues, respectively, in their principal isoforms). The sequences of the CaMKII kinase-hub linkers are consistent with their being intrinsically disordered, allowing the disposition of the kinase domains with respect to the central hub to be variable (Myers et al., 2017).

Our current understanding of the autoregulation of CaMKII is based on studies that have focused primarily on CaMKII-α, and it has been assumed that the other three isoforms function similarly. In this basic scheme, the regulatory segment of CaMKII blocks the kinase active site in the basal state, and this inhibition is released by the binding of Ca^2+^/calmodulin (Ca^2+^/CaM) to the regulatory segment (Figure 1d). The binding of Ca^2+^/CaM facilitates the *trans-* autophosphorylation of Thr 286 within the regulatory segment of one CaMKII subunit by the activated kinase domain of another subunit in the same holoenzyme (Hanson et al., 1994; Rich and Schulman, 1998). How the balance of phosphorylation between the activating and inhibitory sites is controlled in the different isoforms of CaMKII is poorly understood.

We developed a single-molecule assay to measure the phosphorylation status of CaMKII expressed in mammalian cells. In this assay, CaMKII is fused to an N-terminal fluorescent protein (a monomeric variant of enhanced GFP, mEGFP (Cormack et al., 1996; Zacharias et al., 2002)) and to a biotin tag, enabling the capture and visualization, by total internal reflection fluorescence (TIRF) microscopy, of individual CaMKII holoenzymes on a glass slide coated with streptavidin (Figure 2a). Since CaMKII is captured directly from the cell lysate, this procedure avoids difficulties in expressing and purifying various isoforms of CaMKII, particularly those with longer linkers. In addition, the use of a flow cell allows CaMKII holoenzymes to be activated under different reaction conditions after immobilization on glass. Site-specific antibodies that recognize phosphorylated Thr 286 (pThr 286) or phosphorylated Thr 305/Thr 306 (pThr 305/306) can be used to probe the extent of phosphorylation on activating and inhibitory sites on individual holoenzymes (Figure 2a).

**Figure 2:**
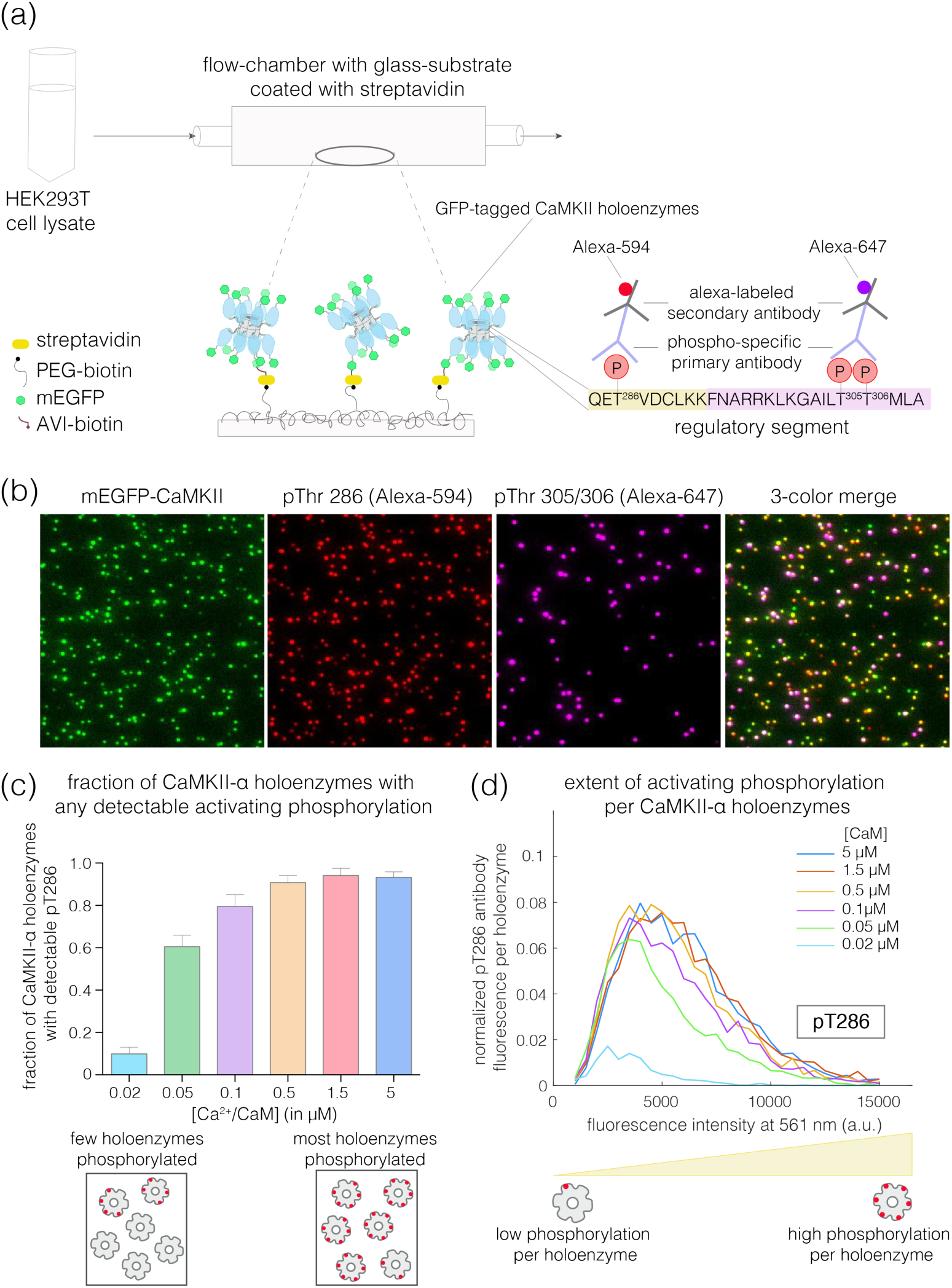
Mammalian expression-based single-molecule Total Internal Reflection Fluorescence (TIRF) assay. (a) Schematic diagram showing the experimental setup. Biotinylated mEGFP-CaMKII overexpressed in HEK 293T cells was pulled down directly from diluted cell lysate, allowing visualization at a single-molecule resolution. The immobilization onto glass substrates functionalized with streptavidin relies on the interaction between biotinylated CaMKII and streptavidin. Autophosphorylation status of CaMKII holoenzyme can be measured using phosphospecific primary antibodies and Alexa-labeled secondary antibodies. (b) Representative single-molecule TIRF images showing mEGFP-CaMKII holoenzymes (green dots), phosphorylation at Thr 286 (red dots) and phosphorylation at Thr 305/306 (purple dots) from left to right. A 3-color merge of these images reports on the fraction of CaMKII holoenzymes that are phosphorylated at Thr 286 and/or Thr 305/306. (c) Fraction of CaMKII-α that shows detectable phosphorylation at Thr 286 is plotted for different Ca^2+^/CaM concentrations ranging from 0.02 μM to 5 μM. The cartoon on the bottom panel depicts two extreme cases, where only a few holoenzymes are phosphorylated or where most holoenzymes are phosphorylated. (d) Distribution of intensity for pThr 286 (561 nm), at different Ca^2+^/CaM concentrations, for CaMKII-α holoenzymes with detectable phosphorylation. The area under each plot is scaled by the fraction of holoenzymes that show no detectable phosphorylation under that condition (see Methods for details of normalization). The cartoon in the bottom panel shows that a right-shift in the peak value of the intensity histogram represents a higher extent of phosphorylation within a CaMKII holoenzyme and *vice versa*.

Using this assay, we discovered that activation of CaMKII-α, which has a 31-residue kinase-hub linker, results in robust autophosphorylation on the activating site (Thr 286), with relatively little inhibitory phosphorylation on Thr 305/306. This is consistent with expectation (Baucum et al., 2015). Unexpectedly, CaMKII-β, with a longer ∼200-residue linker, undergoes robust inhibitory autophosphorylation on Thr 305/306, with less activating phosphorylation on Thr 286. We also monitored autophosphorylation for an artificial construct of CaMKII that is identical to CaMKII-α, except that the kinase-hub linker is replaced by a 218-residue linker from CaMKII-β (Figure 1c). The autophosphorylation pattern for this construct resembles that of CaMKII-β, showing that the difference in activating and inhibitory autophosphorylation between the two isoforms is due entirely to differences in the kinase-hub linkers.

When CaMKII is inactivated after activation, by removing Ca^2+^/CaM and ATP, we found that the dephosphorylation by phosphatases of activating phosphorylation at Thr 286 is much slower than dephosphorylation of inhibitory phosphorylation at Thr 305/306, which is fast. The protection of activating phosphorylation and the rapid reversal of inhibitory phosphorylation by phosphatases allows CaMKII to act as an integrator of Ca^2+^ pulses in cells. CaMKII-α and CaMKII-β can assemble to form mixed holoenzymes, with the balance between activating and inhibitory autophosphorylation determined by the ratio of the two isoforms. Thus, the calcium responsiveness of CaMKII holoenzymes can be tuned by varying the relative levels of the two isoforms.

## Results and Discussion

### A single-molecule assay for measuring CaMKII autophosphorylation

We designed a single-molecule assay to measure autophosphorylation of CaMKII, relying on immediate capture of the holoenzyme after lysis of mammalian cells overexpressing CaMKII (see Methods) (Figure 2a). This procedure is based on a general single-molecule immunofluorescence assay reported previously (Jain et al., 2011). Our assay enables the rapid isolation of CaMKII for single-molecule analysis while minimizing heterogeneity that can result from proteolysis or aggregation during conventional purification. In this assay, GFP-CaMKII holoenzymes, whether or not phosphorylated, are detected with 488 nm laser excitation (green channel). The phosphorylation levels on Thr 286 and Thr 305/306 are monitored using phosphospecific antibodies labeled with Alexa-594 or Alexa-647, detected with 561 nm (red channel) or 640 nm laser excitation (purple channel), respectively (Figure 2a); see Methods. Representative three-color TIRF images from these experiments are shown in Figure 2b.

We quantify the extent of phosphorylation in two ways. First, we measure the fraction of CaMKII holoenzymes that show detectable phosphorylation at Thr 286, regardless of the extent of phosphorylation, by monitoring the co-localization of green and red spots (Figure 2c). To quantify the extent of activating phosphorylation per holoenzyme, we compile intensity distributions for pThr 286 (red channel), taking into account both holoenzymes that are phosphorylated and those that show no detectable phosphorylation (Figure 2d, see Methods for the details of normalization). The integrated intensity of the pThr 286 signal, normalized by the total number of holoenzymes detected (i.e., the number of green spots), reflects the mean level of phosphorylation per holoenzyme (Figure 2d). Phosphorylation at Thr 305/306 is quantified similarly, using the appropriate antibody.

We used this assay to measure autophosphorylation levels for CaMKII upon activation. Human CaMKII-α overexpressed in HEK293T cells and captured from cell lysate is in an inactive state, with no detectable phosphorylation at Thr 286 or Thr 305/306 (data not shown). We activated CaMKII-α by flowing Ca^2+^/CaM and ATP over the glass coverslip to which the captured holoenzymes are tethered (Figure 2a). Increasing the concentration of Ca^2+^/CaM from 20 nM to 5 μM, while maintaining a constant saturating concentration of Ca^2+^ (100 μM), resulted in the detection of increasing levels of phosphorylation at Thr 286 (Figure 2c-d). The EC_50_ value for CaMKII-α activation by Ca^2+^/CaM, derived from these measurements, is in the range of 50-100 nM. This is consistent with previously reported values for the EC_50_ of CaMKII-α activation by Ca^2+^/CaM (Chao et al., 2011; Schulman, 1984; Sloutsky et al., 2019).

A limitation of this assay is that the integrated intensity values cannot be used to derive the absolute number of phosphate groups within each holoenzyme, because we do not know whether every phosphate group necessarily has an antibody bound to it when the maximum signal is obtained. Detection of phosphorylation may be less than complete because of steric interference between antibodies, or interference with the glass support. Due to this limitation, we use the values of the integrated antibody fluorescence intensities to provide a relative, rather than an absolute, measure of the extent of phosphorylation.

### CaMKII-β, with a long kinase-hub linker, acquires inhibitory autophosphorylation more readily than CaMKII-α

We used the single-molecule assay to compare the levels of phosphorylation at the activating (Thr 286) and inhibitory (Thr 305/306) sites in CaMKII-α and CaMKII-β. We expressed CaMKII-α and CaMKII-β separately in HEK293T cells and, in parallel experiments, captured the CaMKII isoforms on glass, followed by activation with high concentrations of Ca^2+^/CaM (5 μM) and 500 μM ATP.

These experiments reveal clear differences in the extent of autophosphorylation at the inhibitory sites in CaMKII-α and CaMKII-β. Whereas 95% of the CaMKII-β holoenzymes show detectable phosphorylation at Thr 305/306, only 30% of the CaMKII-α holoenzymes do so (Figure 3a-right, inset). The mean value of the integrated intensity for Thr 305/306 phosphorylation per holoenzyme is 5-fold higher for CaMKII-β than for CaMKII-α (Figure 3a-right). The α and β isoforms also differ in the extent of activating phosphorylation on Thr 286, with CaMKII-α showing more Thr 286 phosphorylation than CaMKII-β (Figure 3a-left).

**Figure 3:**
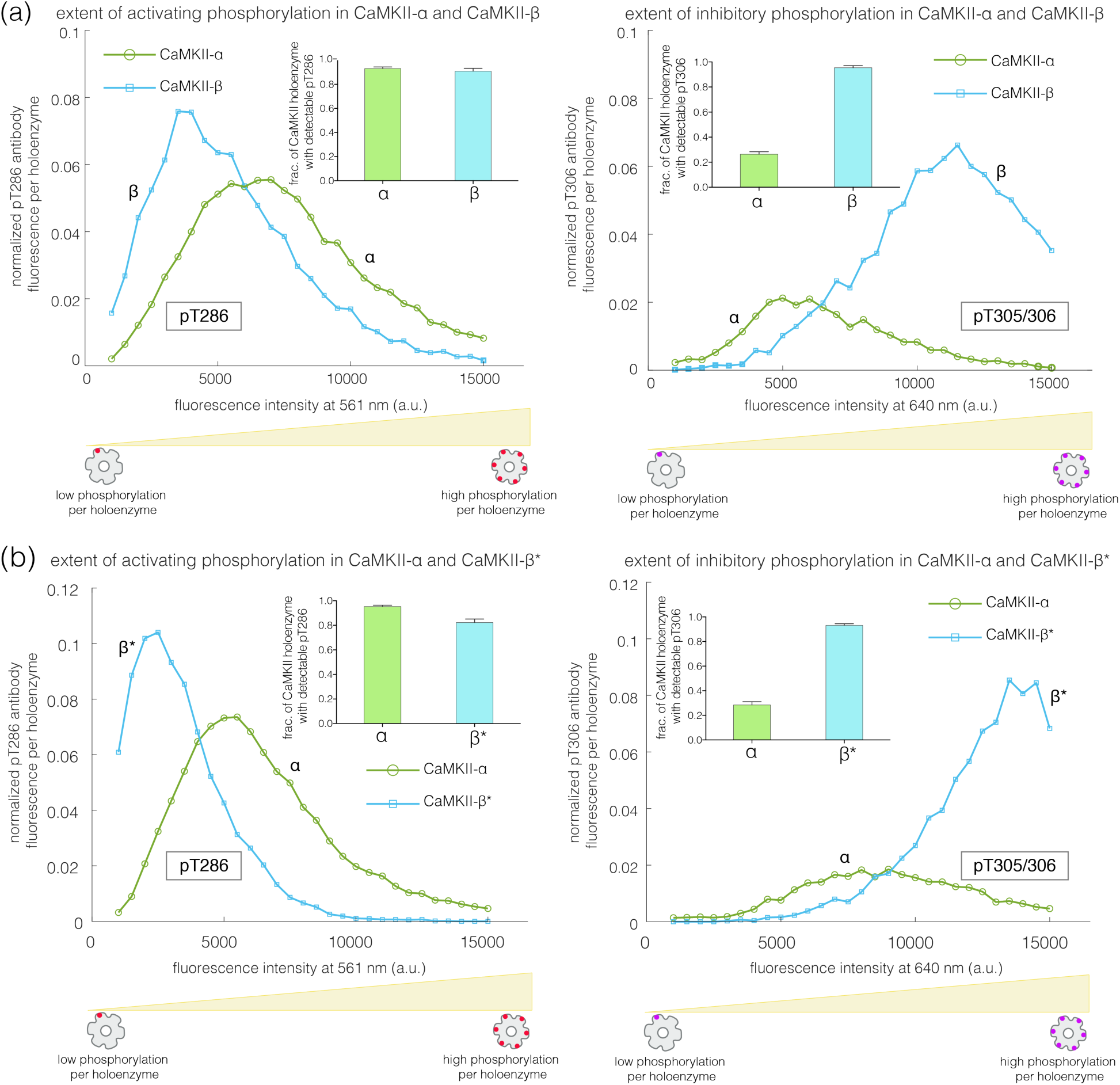
Autophosphorylation status in activated CaMKII-α and CaMKII-β/β*. (a) Comparison of the extent of autophosphorylation (intensity histogram) within CaMKII-α and CaMKII-β holoenzyme at the activating site (Thr 286) (left panel) and the inhibitory site (Thr 305/306) (right panel). The insets show the fraction of holoenzymes that exhibit any detectable phosphorylation for the corresponding phosphosite in CaMKII-α and CaMKII-β. (b) Comparison of the extent of autophosphorylation (intensity histograms) within CaMKII-α and CaMKII-β* holoenzyme at the activating site (Thr 286) (left panel) and the inhibitory site (Thr 305/306) (right panel). The insets show the fraction of holoenzymes that exhibit any detectable phosphorylation for the corresponding phosphosite in CaMKII-α and CaMKII-β*. The area under each intensity histogram is scaled by the fraction of holoenzymes that show no detectable phosphorylation under that condition (see Methods for details of normalization).

We verified that the reduced levels of pThr 305/306 seen for CaMKII-α are not due to an intrinsic inability to phosphorylate this site. We first activated CaMKII-α on glass, as before, and then washed away the Ca^2+^/CaM. This was followed by the addition of ATP to the flow cell. ATP treatment of pre-activated CaMKII led to robust phosphorylation of CaMKII-α on Thr 305/306, at levels comparable to that for CaMKII-β (Figure S3a-c).

We found that the differences in autophosphorylation between the α and β isoforms are due entirely to the differences in the kinase-hub linkers. We created an artificial CaMKII construct in which the kinase-hub linker in CaMKII-α is replaced by that from CaMKII-β (we refer to this construct as CaMKII-β*, because the most distinctive difference between the two isoforms is the nature of the kinase-hub linker, Figure 1c). The autophosphorylation pattern for CaMKII-β* is essentially the same as for CaMKII-β. Each CaMKII-β* holoenzyme exhibits about 5-fold greater phosphorylation of Thr 305/306 than is the case for CaMKII-α (Figure 3b-right), with about 95% of the CaMKII-β* holoenzymes showing detectable phosphorylation at pThr 305/306. CaMKII-β* also shows a lower degree of activating phosphorylation on Thr 286 when compared to CaMKII-α (Figure 3b-left).

We use CaMKII-β* as a surrogate for CaMKII-β in most of the experiments reported in this paper. The sequences flanking the phosphosites being probed by the antibodies are the same in CaMKII-β* and CaMKII-α, whereas there are a few small differences in sequence between CaMKII-β and CaMKII-α in these regions. Thus, when comparing CaMKII-β* to CaMKII-α, observed differences in autophosphorylation are not due to differences in the affinities of the antibodies for their cognate sites.

CaMKII-α and CaMKII-β can assemble into mixed holoenzymes containing both isoforms (Bennett et al., 1983; Brocke et al., 1999; Thiagarajan et al., 2002). We co-transfected mEGFP-CaMKII-α and mCherry-CaMKII-β* in HEK293T cells in four different ratios (1:0, 3:1, 1:1, and 0:1, α:β*, respectively) and activated the resultant holoenzymes after immobilization on glass (Figure 4a). The co-transfection of mEGFP-CaMKII-α (488 nm) and mCherry-CaMKII-β* (561 nm) resulted in predominantly heterooligomeric holoenzymes, with ∼80% colocalization of red and green spots (data not shown). As the proportion of CaMKII-β* increases, there is an increase in the extent of Thr 305/306 phosphorylation (Figure 4b-c). Thus, alterations in the relative levels of CaMKII-α and CaMKII-β can result in the generation of mixed CaMKII holoenzymes with altered responsiveness to Ca^2+^/calmodulin.

**Figure 4:**
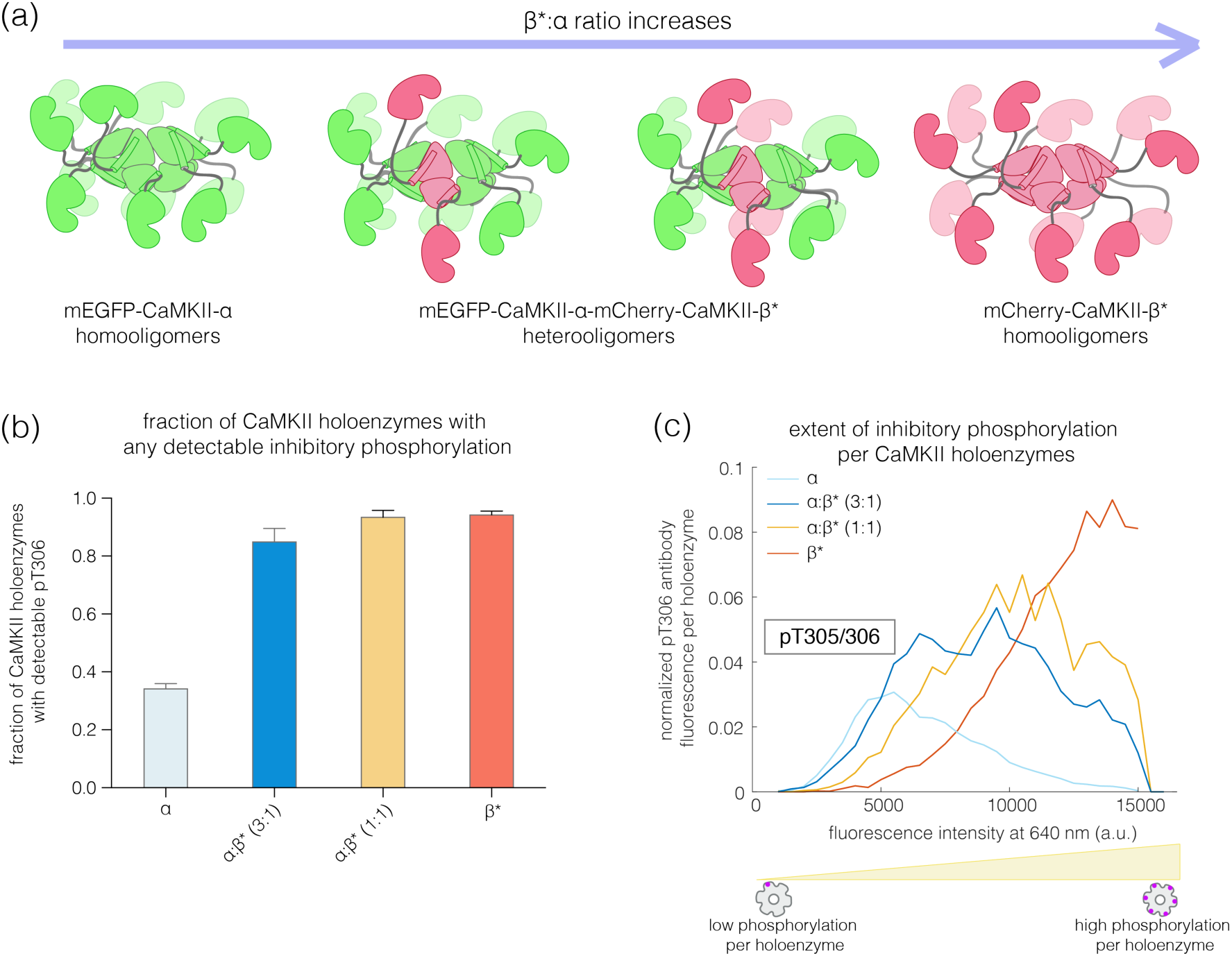
Inhibitory autophosphorylation in CaMKII-α-CaMKII-β* heterooligomers. (a) Schematic diagram showing that co-expression of GFP-CaMKII-α and mCherry-CaMKII-β* leads to the formation of heterooligomers. (b) Bar graph showing the fraction of holoenzymes that show detectable phosphorylation at the inhibitory site (Thr 305/306), which increases as the ratio of CaMKII-β* increases. (c) Intensity histogram for the homooligomers and heterooligomers. As the ratio of CaMKII-β* increases, there is a right-shift in the peak value of the intensity histogram. The area under each intensity histogram is scaled by the fraction of holoenzymes that show no detectable phosphorylation under that condition (see Methods for details of normalization).

### The altered balance of phosphorylation in CaMKII-α and CaMKII-β may arise due to differences in the rates of *cis* and *trans* autophosphorylation in the two isoforms

To understand how changes in the length of the kinase-hub linker can alter the balance of autophosphorylation, we analyzed a simple kinetic model for CaMKII activation in which the holoenzyme contains only two kinase domains (see Appendix). A kinetic model for the activation of a dodecameric CaMKII holoenzyme requires the specification of an extremely large number of intermediate states, and we have not pursued this. In our simple kinetic model, we assume that Thr 286 can only be phosphorylated in *trans*, because it is located too far from the active site of the kinase (Figure 1b, and see Figure 5). We assume that Thr 305/306 can either be phosphorylated *in cis*, as suggested by crystal structures (Rellos et al., 2010), or *in trans*.

**Figure 5:**
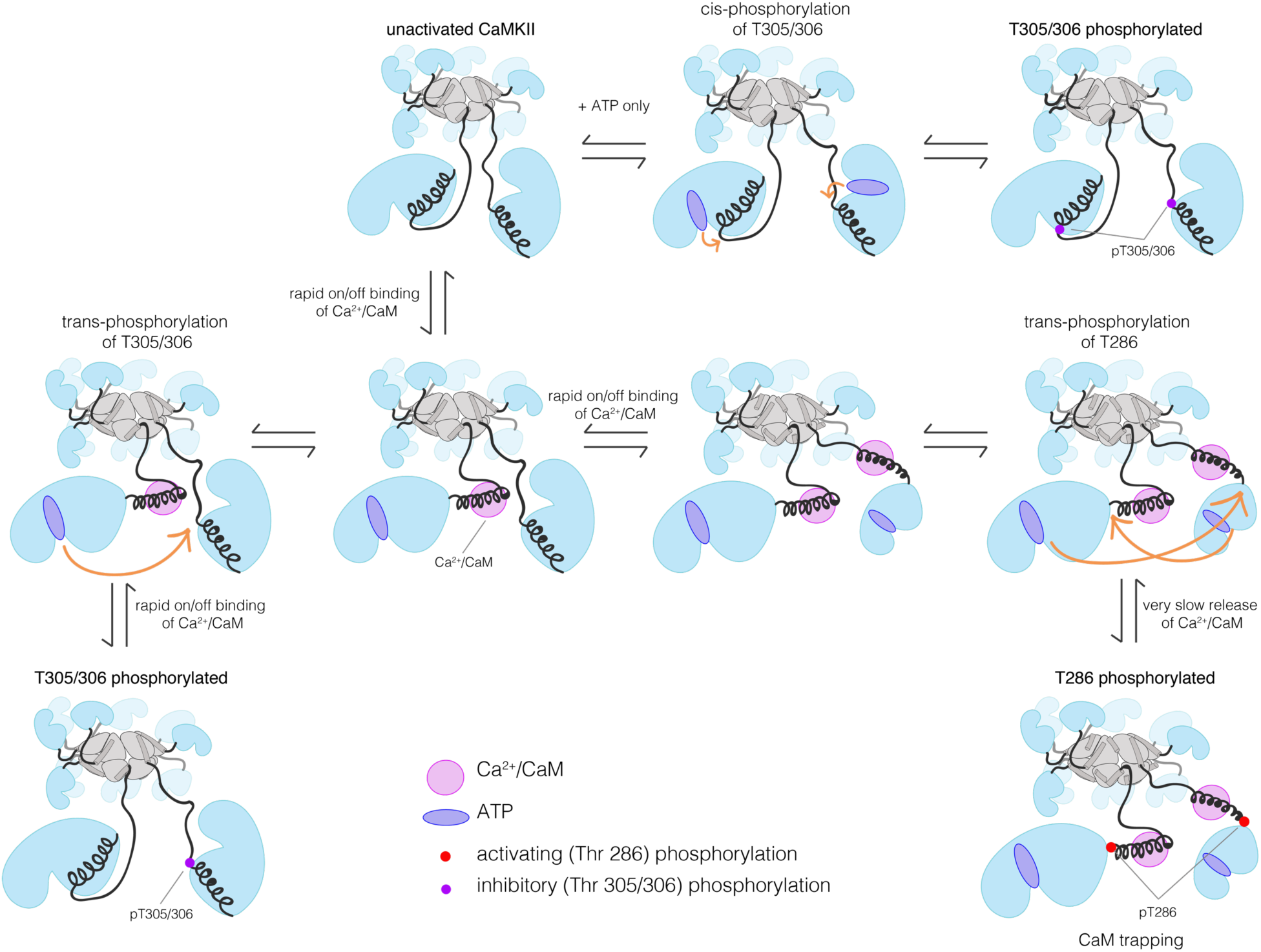
Schematic diagram showing the different modes of autophosphorylation at the activating and inhibitory sites, in the absence or presence of Ca^2+^/CaM. While Thr 305/306 can get phosphorylated both in *cis* in the absence of Ca^2+^/CaM or in *trans* in the presence of Ca^2+^/CaM, autophosphorylation of Thr 286 can only happen in *trans* in the presence of Ca^2+^/CaM. Ca^2+^/CaM shows a rapid association and dissociation until CaMKII gets phosphorylated at Thr 286, when the affinity for Ca^2+^/CaM increases, with a 1000-fold slow release rate. These different reactions and conditions form the basis of our kinetic model.

For *trans*-phosphorylation of either Thr 286 or Thr 305/306, the kinase acting as the enzyme has to have Ca^2+^/CaM bound to it, otherwise the active site of the enzyme will be blocked. For *trans*-phosphorylation of Thr 286, the kinase acting as the substrate must also have Ca^2+^/CaM bound to it, otherwise the Thr 286 is not accessible for phosphorylation (Rich and Schulman, 1998). Autophosphorylation of Thr 305/306 can only occur in the absence of Ca^2+^/CaM binding to the substrate kinase, because otherwise Thr 305 and Thr 306 would be covered by Ca^2+^/CaM. A key idea behind this model is that we assume that the longer linker in CaMKII-β slows down the rates of *trans*-phosphorylation reactions within a holoenzyme when compared to CaMKII-α, but that the rates of *cis*-phosphorylation are the same as in the two isoforms (Sørensen and Kjaergaard, 2019).

We analyzed the predicted autophosphorylation kinetics for the CaMKII dimer model using reasonable estimates for the rate constants in the model (see Appendix). Our results show that when the rate of *trans*-phosphorylation decreases relative to the rate of *cis*-phosphorylation, as is expected to occur with longer kinase-hub linkers, then there is a switch from Thr 286 phosphorylation dominating to Thr 305/306 phosphorylation dominating (Figure S5a-b, see Appendix). If Thr 286 is phosphorylated, the affinity for Ca^2+^/CaM increases ∼1000 fold, a phenomenon referred to as the calmodulin-trapping (Meyer et al., 1992; Tse et al., 2007). Since Ca^2+^/CaM covers the inhibitory site (Thr 305/306), phosphorylation at this site is suppressed (Figure 5). The situation is reversed when the linker is long, and *cis*-phosphorylation is faster than *trans*-phosphorylation (Figure S5b). In this case, activating phosphorylation is suppressed, because Ca^2+^/CaM binding is blocked.

### The balance between inhibitory and activating autophosphorylation is maintained upon activation of CaMKII in cells

We tested whether the balance between activating and inhibitory autophosphorylation is maintained when the enzyme activated in cells. We overexpressed CaMKII-α or CaMKII-β* in HEK293T cells, along with calmodulin, followed by ionomycin treatment to increase intracellular Ca^2+^ levels. The cells were then lysed and CaMKII holoenzymes were captured and assayed as before.

Phosphorylation at Thr 286 increases upon treatment with ionomycin for both CaMKII-α and CaMKII-β*, as expected (Baucum et al., 2015) (Figure S3d). CaMKII-α shows higher phosphorylation at Thr 286 as compared to CaMKII-β*, in agreement with our *in vitro* results (Figure S3d). However, no phosphorylation at Thr 305/306 is detected for either CaMKII-α or CaMKII-β* in this experiment (Figure S3e), which is consistent with previous reports (Baucum et al., 2015). There is depletion of ATP levels upon cell lysis, which could result in reduced kinase activity in the presence of phosphatase activity. We therefore incubated cells with okadaic acid and cyclosporine A (250 nM each), which are inhibitors of PP1 and calcineurin, respectively, for 45-60 minutes before activation. This treatment led to detectable levels of phosphorylation at Thr 305/306 for ∼30% of the CaMKII-β* holoenzymes, but not for CaMKII-α (Figure S3e). This is consistent with the increased levels of Thr 305/306 phosphorylation seen for CaMKII-β* using the single-molecule assay.

### Activating autophosphorylation is more resistant to dephosphorylation, compared to inhibitory phosphorylation

We incubated CaMKII-α and CaMKII-β* with Ca^2+^/CaM and ATP on glass, while including increasing amounts of λ-phosphatase, a potent non-specific phosphatase (Zhuo et al., 1993), in the activation buffer (Figure 6a). Active CaMKII-α is efficient at maintaining phosphorylation on Thr 286 in the face of competing phosphatase activity, while CaMKII-β* is less so (Figure 6b, d). For CaMKII-β*, high levels of inhibitory phosphorylation at Thr 305/306 are maintained in the presence of phosphatase (Figure 6c, e). In contrast, phosphorylation at Thr 305/306 is completely removed by the phosphatase for CaMKII-α (Figure 6c, e). Thus, the conclusion that CaMKII-α and CaMKII-β/β* more readily gain activating and inhibitory phosphorylation, respectively, holds true in the presence of phosphatases during activation.

**Figure 6:**
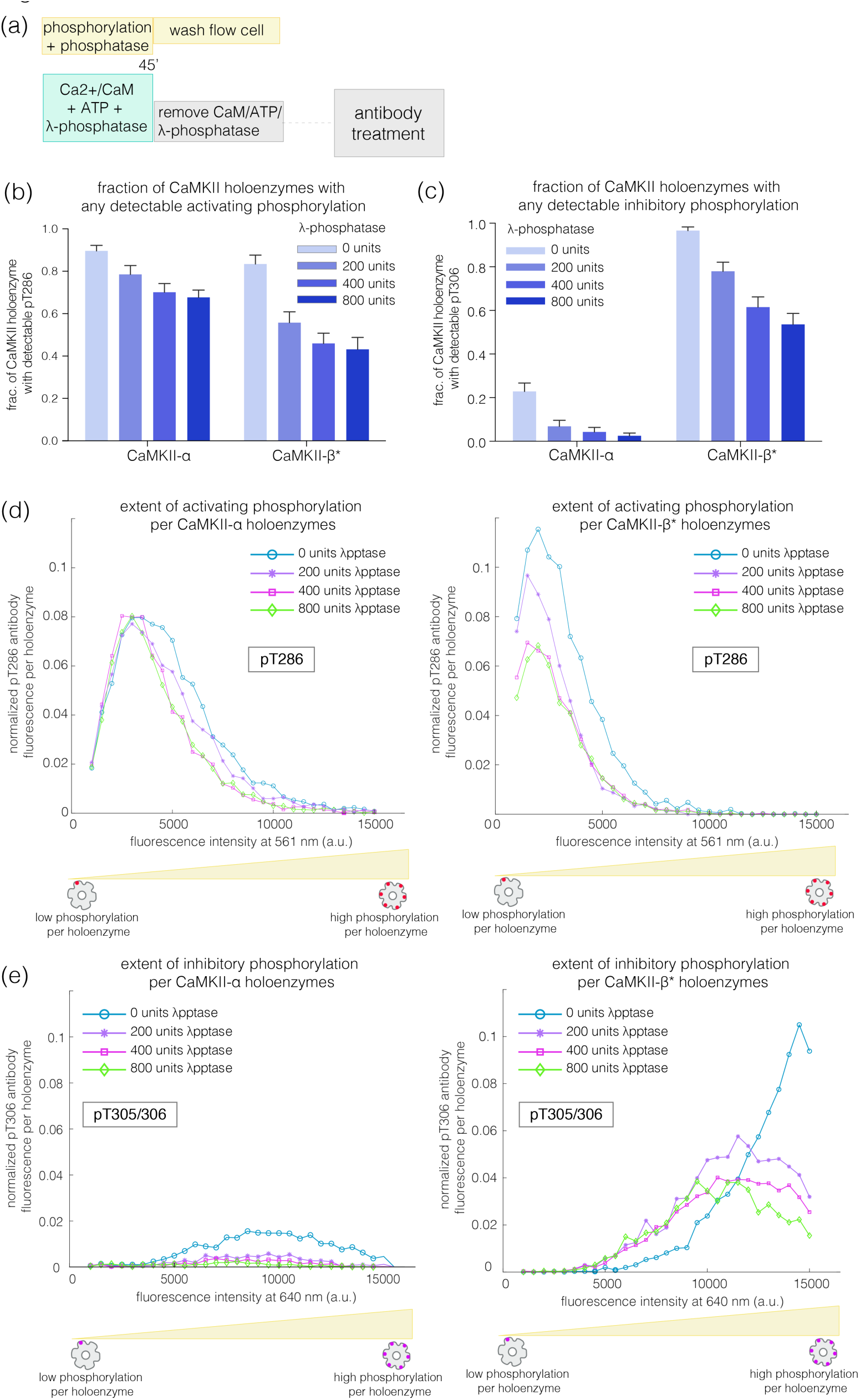
Effect of λ-phosphatase on the phosphorylation status of CaMKII when the kinase is active. (a) Schematic diagram showing the experimental set up. CaMKII was activated in the presence of 0, 200, 400, or 800 units of λ-phosphatase for 45 min. (b-c) Bar graph showing the fraction of CaMKII-α and CaMKII-β* holoenzymes that shows detectable phosphorylation at the activating site (Thr 286) and the inhibitory site (Thr 305/306), respectively, in the presence of λ-phosphatase. (d) Intensity distribution of pThr 286 (561 nm) signal for CaMKII-α (left panel) and CaMKII-β* (right panel) holoenzymes with detectable phosphorylation in the presence of λ-phosphatase. (e) Intensity distribution of pThr 305/306 (640 nm) signal for CaMKII-α (left panel) and CaMKII-β* (right panel) holoenzymes with detectable phosphorylation in the presence of λ-phosphatase. The area under each plot in (d-e) is scaled by the fraction of holoenzymes that show no detectable phosphorylation under that condition (see Methods for details of normalization).

We next investigated how efficiently the phosphatase can dephosphorylate the activating and inhibitory sites when the kinase activity is switched off, as would happen after a calcium pulse. We first activated CaMKII on glass, and then washed away Ca^2+^/CaM and ATP (see Methods, Figure 7a). In the previous experiments, we determined the concentration of phosphatase that results in the maximum reduction of phosphorylation levels during activation – we refer to this concentration of phosphatase as a “saturating” concentration. With the kinase activity switched off, we initiated dephosphorylation reactions by adding a saturating concentration of λ-phosphatase to the phosphorylated CaMKII. The dephosphorylation reaction was then stopped by washing away the phosphatase at defined time points. The extent of phosphorylation at Thr 286 and Thr 305/306 was then measured as before (Figure 7a).

**Figure 7:**
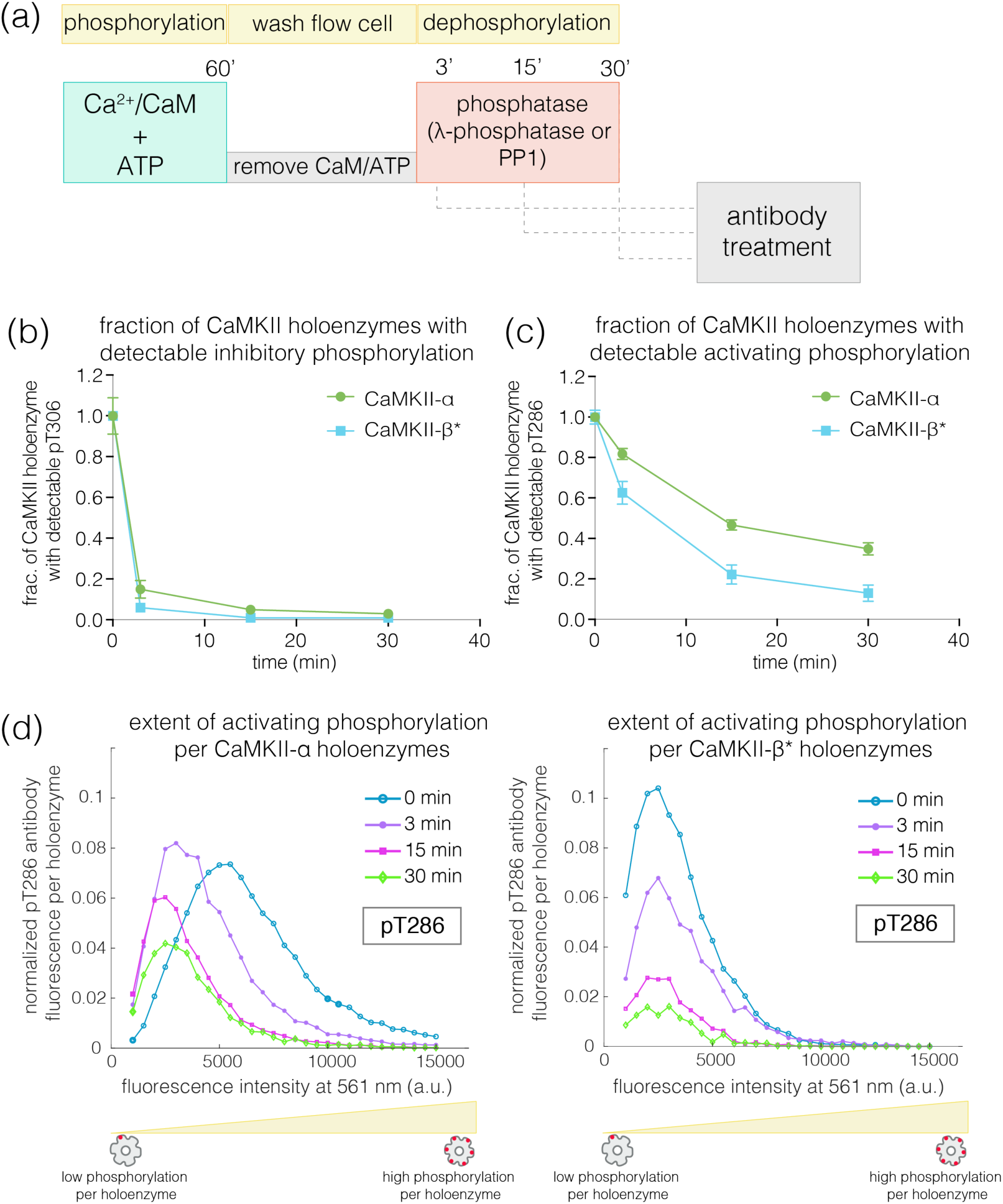
Effect of λ-phosphatase on the dephosphorylation kinetics when kinase activity is switched off. (a) Schematic diagram showing the experimental set up. CaMKII was activated, followed by a wash to remove the components of the activation buffer, and then saturating amounts of λ-phosphatase/PP1α (400-800 units) were added for 0, 3, 15, or 30 min. (b) Plot showing the fraction of α and β* holoenzymes that exhibits detectable phosphorylation at the inhibitory site (Thr 305/306) upon treatment with λ-phosphatase for defined time-points. The fractions at 3, 15, or 30 minutes for α and β* are normalized by the corresponding activated version that has not been exposed to any λ-phosphatase (0-minute time-point, whose value is set to 1.0). (c) Same as (b) but for the activating site (Thr 286). (d) Intensity distribution for pThr 286 (561 nm) for CaMKII-α (left panel) and CaMKII-β* (right panel) holoenzymes with detectable phosphorylation after 0, 3, 15, or 30 minutes of λ-phosphatase treatment. The area under each plot is scaled by the fraction of holoenzymes that show no detectable phosphorylation under that condition (see Methods for details of normalization).

In the absence of competing kinase activity, pThr 305/306 is rapidly dephosphorylated, for both CaMKII-α and CaMKII-β* (Figure 7b, Figure S7a). In contrast, the rate of dephosphorylation of pThr 286 is slow, with complete dephosphorylation taking more than 30 minutes for both isoforms (Figure 7c-d, Figure S7a). We also measured the dephosphorylation kinetics of CaMKII-α and CaMKII-β* using the catalytic subunit of protein phosphatase 1α (PP1α), an endogenous phosphatase for CaMKII. Slower rates of dephosphorylation at pThr 286 and rapid dephosphorylation at pThr 305/306 are observed for CaMKII-α and CaMKII-β* using PP1α, as for λ-phosphatase (Figure S7b-c).

To rule out that the resistance of pThr 286 towards rapid dephosphorylation is a consequence of on-glass treatments, we performed experiments in solution using diluted HEK293T cell lysate (see Methods) (Figure S7d). CaMKII-α and CaMKII-β* were first activated in the diluted cell lysate and then treated with a high concentration of staurosporine (100 μM), which inhibits kinase activity (Figure S7g). We then inactivated calmodulin by chelating Ca^2+^ by EGTA (1 mM), followed by the addition of saturating amounts of λ-phosphatase. Samples were pulled down from this reaction mixture at defined time points, and the phosphorylation at Thr 286 and Thr 305/306 was measured using the single-molecule assay. These experiments also showed slower rates of dephosphorylation at Thr 286 when compared to Thr 305/306 (Figure S7e-f).

The observed resistance of pThr 286 towards rapid dephosphorylation suggests that this site may be less accessible to phosphatases compared to pThr 305/306. Unexpectedly, we found that the addition of Ca^2+^/CaM to phosphorylated CaMKII enhances the rate of dephosphorylation at Thr 286 by about 4-fold for both CaMKII-α and CaMKII-β* (Figure 8a-c, Figure S8). When Ca^2+^/CaM is present, the pThr 286 signal is reduced at a rate that is comparable to the dephosphorylation rate of pThr 305/306. This effect does not require the addition of ATP, i.e., it is independent of kinase activity. The slower rates of dephosphorylation at Thr 286 is also noted for CaMKII-β, with the addition of Ca^2+^/CaM reversing the protection (data not shown).

**Figure 8:**
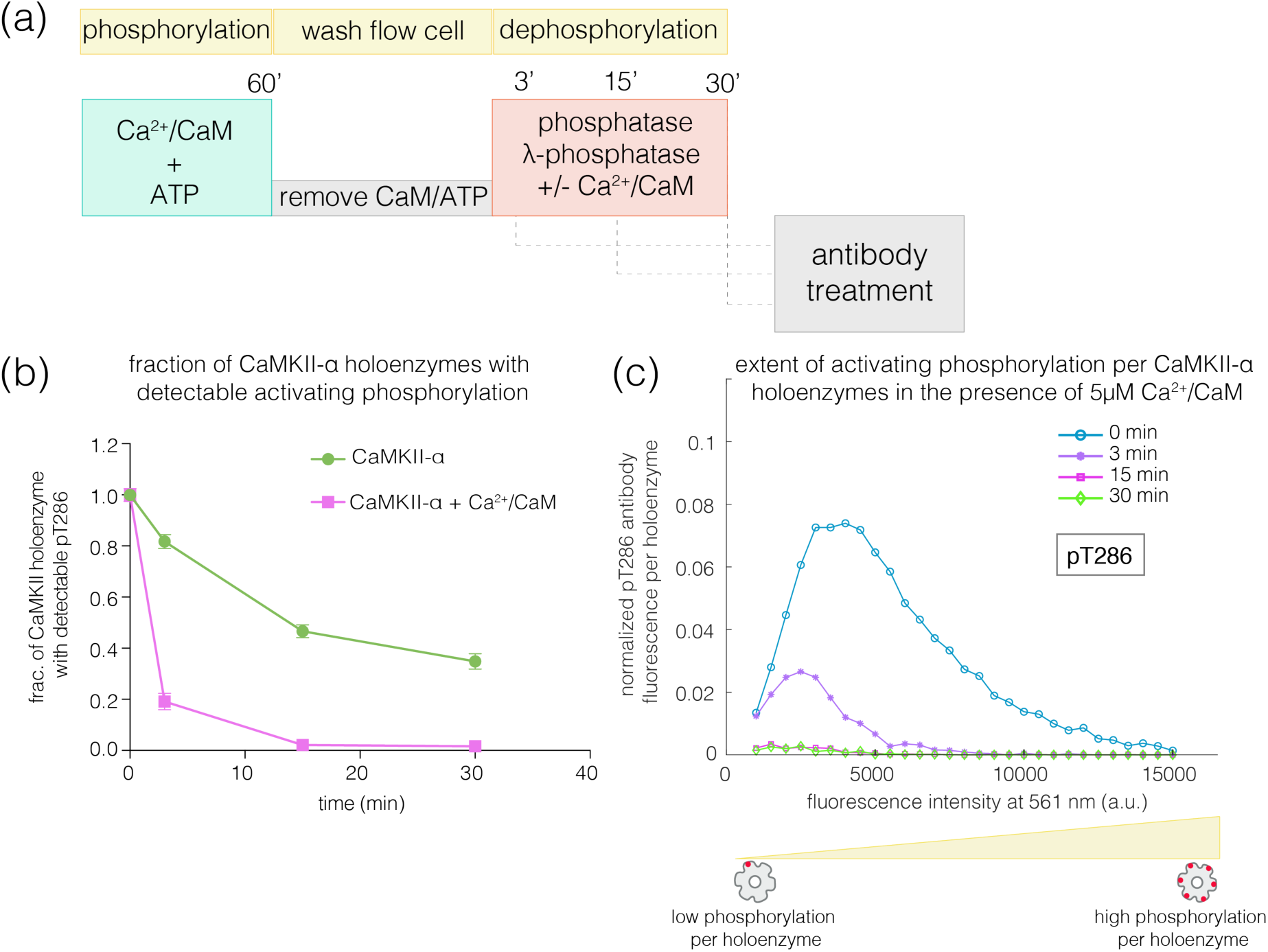
Effect of addition of Ca^2+^/CaM on the rates of dephosphorylation at the activating site. (a) Schematic diagram showing the experimental set up. CaMKII was activated, followed by wash to remove the components of the activation buffer and then saturating amounts of λ-phosphatase were added for 3, 15, or 30 min, in the presence and absence of Ca^2+^/CaM. (b) Plot showing the fraction of CaMKII-α holoenzymes with detectable phosphorylation at the activating site (Thr 286, right panel) after 0 minute (activated control) and 3, 15, or 30 minutes of treatment with saturating amounts of λ-phosphatase in the absence (green trace) and presence (pink trace) of Ca^2+^/CaM. The fractions at 3, 15, or 30 minutes are normalized with respect to activated CaMKII that has not been exposed to any λ-phosphatase (0-minute, whose value is set to 1.0). (c) Intensity distribution for pThr 286 (561 nm) for CaMKII-α holoenzymes with detectable phosphorylation, upon 0, 3, 15, or 30 minutes of λ-phosphatase treatment in the presence of 5 μM Ca^2+^/CaM. The area under each plot is scaled by the fraction of holoenzymes that show no detectable phosphorylation under that condition (see Methods for details of normalization).

The addition of Ca^2+^/CaM has no effect on the rate of dephosphorylation of pThr 305/306, indicating that Ca^2+^/CaM does not increase the activity of the phosphatase. These data suggest that, in the absence of Ca^2+^/CaM, the calmodulin-binding element of CaMKII plays a role in sequestering pThr 286 from phosphatases. The mechanism underlying this protection remains to be understood.

### Slower dephosphorylation at the activating site primes CaMKII for activation by incoming Ca^2+^ pulses

The slower rates of dephosphorylation at Thr 286 might ensure that at least some subunits in a CaMKII holoenzyme retain Ca^2+^/CaM-independent activity after a calcium pulse has subsided. CaMKII subunits that are phosphorylated on Thr 286 have increased affinity for Ca^2+^/CaM, which can allow a subthreshold Ca^2+^-pulse to cooperatively spread Thr 286 phosphorylation within the primed holoenzyme (De Koninck and Schulman, 1998; Meyer et al., 1992; Tse et al., 2007). We tested this by generating a population of CaMKII holoenzymes in which only a small number of subunits are phosphorylated at Thr 286, to determine if they are primed for a response to Ca^2+^/CaM.

Treatment of activated CaMKII-α with λ-phosphatase for 3-5 minutes generated a population in which about 50% of the holoenzymes retained some phosphorylation at Thr 286, while the rest of the holoenzymes had no detectable phosphorylation remaining (Figure 9a-c). The holoenzymes that do retain Thr 286 phosphorylation have reduced levels of phosphorylation per holoenzyme (Figure 9d). When such a sample is incubated with ATP and a sub-saturating level of Ca^2+^/CaM (25 nM), the extent of phosphorylation for holoenzymes that show detectable pThr 286 is restored to the levels seen for fully activated CaMKII-α (Figure 9a-b, d). Treatment with ATP and sub-saturating Ca^2+^/CaM does not increase the fraction of CaMKII-α holoenzymes with detectable phosphorylation at Thr 286 very much, suggesting that some extent of pre-existing phosphorylation on Thr 286 is important for gaining additional phosphorylation with sub-saturating levels of Ca^2+^/CaM (Figure 9c).

**Figure 9:**
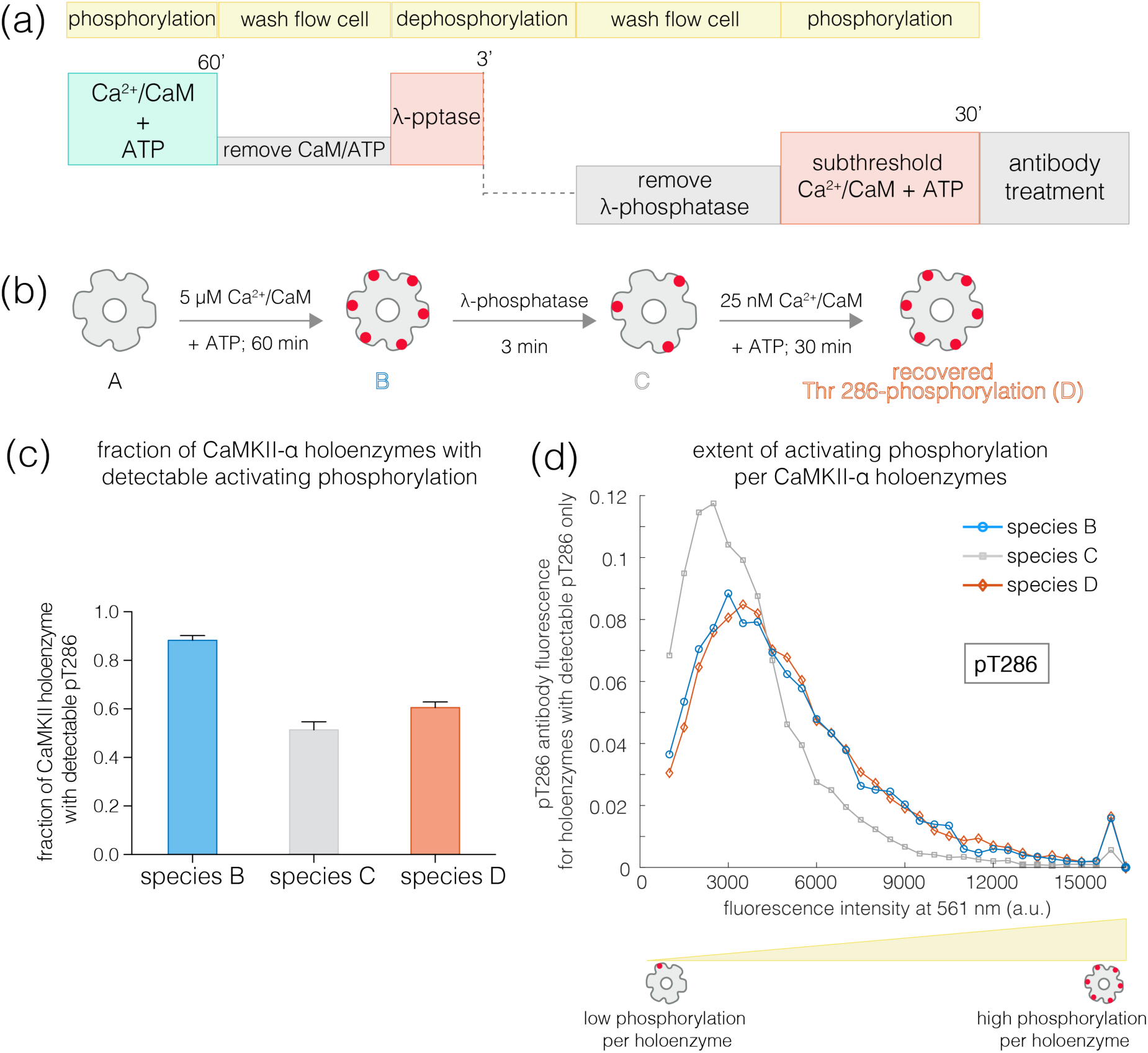
Recovery of phosphorylation at the activating site by subthreshold concentrations of Ca^2+^/CaM. (a) Schematic diagram showing the experimental set up. CaMKII was activated, followed by a wash to remove the components of the activation buffer and then saturating amounts of λ-phosphatase were added for 3-5 min. The sample is then washed to remove the λ-phosphatase, followed by further treatment with subthreshold concentrations of Ca^2+^/CaM (25 nM) for 30 minutes and the autophosphorylation status was measured. (b) Cartoon representation of the different species generated after each treatment. Each species is color-coded and the color schemes are maintained throughout the plots. (c) Fraction of CaMKII-α holoenzymes with detectable phosphorylation at Thr 286 is plotted for each species. (d) Intensity distribution of pThr 286 (561 nm) for only those CaMKII-α holoenzymes that show any detectable Thr 286 phosphorylation, for the different species of interest as described in (b).

A similar experiment using CaMKII-β* yields very different results. Here, phosphatase treatment results in substantial reduction in pThr 286 levels. Subsequent treatment with 25 nM Ca^2+^/CaM and ATP does not result in a marked increase in the extent of Thr 286 phosphorylation per holoenzyme. In contrast, ∼40% of CaMKII-β* holoenzymes show detectable phosphorylation at Thr 305/306 after treatment with sub-saturating Ca^2+^/CaM, compared to ∼2% before (data not shown). Thus, in contrast to CaMKII-α holoenzymes, CaMKII-β holoenzymes tend to become inactivated, rather than undergoing priming for further activation.

### Concluding Remarks

CaMKII is unusual among protein kinases because it is assembled into a large holoenzyme with twelve to fourteen subunits. Analysis of the phosphorylation status of the enzyme using bulk assays, such as Western blots, has been challenging due to difficulties in purifying the holoenzymes, particularly for isoforms with longer kinase-hub linkers. In addition, bulk measurements do not provide a window into the extent of phosphorylation per holoenzyme. To overcome these limitations, we designed a single-molecule TIRF microscopy assay that relies on the capture from mammalian cell lysates of CaMKII holoenzymes tagged with fluorescent proteins. Immobilization of the captured holoenzymes in a flow-cell apparatus allows activation and dephosphorylation of CaMKII to be carried out readily, followed by measurement of the phosphorylation status using fluorescently labeled site-specific antibodies. Using this assay, we discovered that CaMKII-α and CaMKII-β, the two major isoforms in the brain, differ in the balance of activating and inhibitory phosphorylation. CaMKII-α, with a shorter linker, readily acquires activating phosphorylation, while CaMKII-β, with a longer linker, is biased towards inhibitory phosphorylation.

The difference in autophosphorylation balance between CaMKII-α and CaMKII-β can be explained by the fact that Thr 286 can only be autophosphorylated in *trans*, between two kinase domains, since this residue is located too far from the active site of the subunit of which it is a part. Phosphorylation of Thr 286 is known to occur predominantly through *trans*-autophosphorylation within a holoenzyme, rather than phosphorylation between different holoenzymes (Rich and Schulman, 1998). In contrast, inhibitory phosphorylation on Thr 305/306 can occur either *in cis* or *in trans*. We expect that the longer linker length in CaMKII-β results in a reduction in the rate of *trans*-autophosphorylation of Thr 286, while the rate of *cis*-autophosphorylation of Thr 305/306 is the same for CaMKII-α and CaMKII-β. As the kinase-hub linker length increases, the rate of activating *trans*-phosphorylation decreases, which switches the balance towards inhibitory phosphorylation in CaMKII-β. The switch is expected to be sharpened by the fact that inhibitory autophosphorylation blocks Ca^2+^/CaM binding, thereby preventing activating phosphorylation.

There is a parallel between the architecture of CaMKII and A-kinase anchoring proteins (AKAPs), which are connected to cAMP-dependent protein kinase (PKA) by long flexible linkers. It has been suggested that the length and flexibility of the AKAP linker determines the spatial range over which PKA can phosphorylate its substrates (Smith et al., 2013). Since CaMKII holoenzymes are anchored within the post-synaptic density, the differences in the lengths of the kinase-hub linkers in different isoforms may likewise control the spatial extent of substrate phosphorylation enabled by the activation of a particular CaMKII holoenzyme. Our work has now revealed a key difference between CaMKII and AKAP:PKA complexes, which is that the kinase-hub linker length in CaMKII can control the activation status of the holoenzyme, not just the spatial range of the kinase.

Our experiments show that activating phosphorylation on Thr 286 is resistant to dephosphorylation, whereas inhibitory phosphorylation on Thr 305/306 is reversed rapidly. This suggests that the phosphate group on Thr 286 is somehow protected from phosphatases. The protection afforded to pThr 286 is released by the addition of Ca^2+^/CaM, suggesting that the calmodulin-binding element of CaMKII adopts a conformation that protects pThr 286 in the absence of Ca^2+^/CaM. The presence of such a conformational change may explain the apparent discrepancy between our results and studies reporting that pThr 286 undergoes very rapid dephosphorylation in neurons (Lee et al., 2009). The conclusion that pThr 286 is rapidly dephosphorylated relied on the utilization of a FRET-based sensor, Camui (Takao et al., 2005), which reports on conformational changes undergone by CaMKII, and does not report on phosphorylation directly. It will be interesting to see whether the release of Ca^2+^/CaM from CaMKII after activation results in a compaction of CaMKII that might explain both the protection of pThr 286 that we observe and the rapid relaxation back to a compact state that is reported by the Camui sensor.

The observed resistance of pThr 286 to rapid dephosphorylation by phosphatases can enable CaMKII activity to persist for some time after the decay of Ca^2+^ pulses. In addition, CaMKII subunits that are phosphorylated on Thr 286 have higher affinity for Ca^2+^/CaM, which can facilitate the rebinding of Ca^2+^/CaM during subsequent pulses of calcium. Due to the cooperativity of Ca^2+^/CaM binding to CaMKII, such a priming mechanism enables CaMKII holoenzymes that have retained some activating phosphorylation to rapidly reacquire higher levels of phosphorylation. This may enable the history of synaptic activity, including the frequency and the gaps between periods of activity, to be encoded in CaMKII.

There are cellular differences in the relative levels of CaMKII-α and CaMKII-β and these can be regulated by synaptic activity (Thiagarajan et al., 2002). Their relative ratios have been shown to express as differences in their cellular targeting and stimulus frequency responses, and an important difference described here are their distinct activating and inhibitory autophosphorylation properties, regulated by kinase-hub linker length. The lower inhibitory autophosphorylation of CaMKII-α relative to that in CaMKII-β can have a number of consequences. Autophosphorylation of the inhibitory site reduces binding of CaMKII-α to the postsynaptic density (PSD) *in vitro* (Strack et al., 1997). By contrast, abrogation of such autophosphorylation greatly reduces dissociation of the kinase from PSD in neurons (Shen and Meyer, 1999) and increases PSD binding in transgenic animals, thereby altering synaptic plasticity and long-term potentiation (Elgersma et al., 2002). The rapid inhibitory phosphorylation of CaMKII-β, on the other hand, limits binding of Ca^2+^/CaM, which is required for dissociation of the kinase from F-actin (Shen and Meyer, 1999; Wang et al., 2019), an important step in structural remodeling of the synapse (Kim et al., 2015). Our results show that by varying the relative expression levels of α and β-isoforms, the cell can generate CaMKII heterooligomers with variable response to activating signals, thereby fine-tuning the response of CaMKII to Ca^2+^ spike trains in the brain.

### Appendix: A kinetic model to explain the switch between activating and inhibitory autophosphorylation tendencies between CaMKII-α and CaMKII-β

A simple kinetic model is built to explain the observed differences in the balance between activating and inhibitory autophosphorylation. The model is based on a CaMKII dimer, which avoids the extremely large number of intermediate states that need to be considered for a CaMKII dodecamer. Phosphorylation at the inhibitory site (Thr 305/306) can occur both in *cis* and in *trans*, whereas Thr 286 can only be phosphorylated in *trans*.

We wrote a series of chemical reactions that can lead to autophosphorylation at Thr 286 and Thr 305/306. A representative set of values for the forward and reverse rate constants are shown below. The reaction kinetics corresponding to these equations were simulated using Berkeley Madonna, a general-purpose differential equation solver (http://www.berkeleymadonna.com/). We monitored the accumulation of all species that are phosphorylated at either Thr 286 or Thr 305/306 under different rates of *trans*-phosphorylation, keeping the rates of *cis*-phosphorylation constant (Figure S5a-b).

The on and off rates for Ca^2+^/CaM are set to 10^5^ M^-1^ sec^-1^ (denoted onrate) and 10 sec^-1^ (denoted offrate) respectively, corresponding to a dissociation constant of 10^-4^ M^-1^ prior to CaM trapping. We mimicked the effect of CaM trapping by ignoring the dissociation of Ca^2+^/CaM from species that are phosphorylated at Thr 286. We ignore the effects of phosphatases in this simulation, and set the rate of dephosphorylation to be negligible (10^-9^ sec^-1^, denoted reverserate). The initial concentrations of CaM and CaMKII are set to 10^-4^ M and 10^-9^ M, respectively. The following equations are considered, where K denotes the kinase (CaMKII) and C denotes CaM. Phosphorylation on Thr 286 or Thr 305/306 is denoted by 286 and 306, respectively.

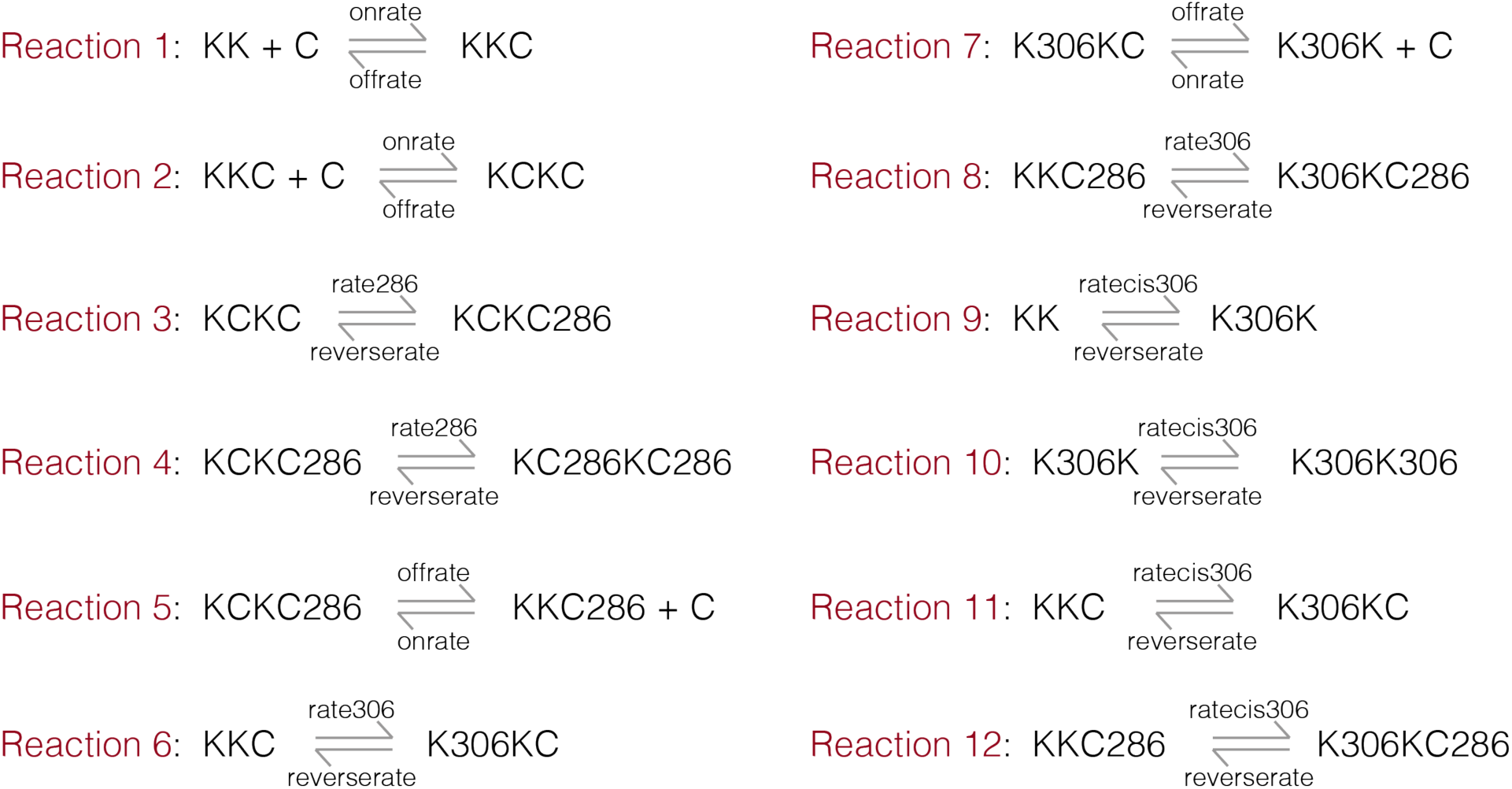

We ran the kinetic simulations for two sets of parameters. For the first set, the value of *k*_cat_ for the kinase reaction is set to 1 sec^-1^ and 0.1 sec^-1^ for the *trans*-phosphorylation of Thr 286 (denoted rate286) and Thr 305/306 (denoted rate306), respectively. The rate of *cis*-phosphorylation of Thr 305/306 (denoted ratecis306) is set to 0.1 sec^-1^. Under these conditions, the simulation shows a dominant accumulation of species that are phosphorylated on Thr 286 (Figure S5a). Next, to mimic the effect of increasing the kinase-hub linker length, the rate of *trans*-phosphorylation was reduced by 10-fold for both Thr 286 and Thr 305/306 (i.e., 0.1 sec^-1^ and 0.01 sec^-1^, respectively), while keeping the rate of *cis*-phosphorylation unaltered. Under these conditions, the simulations show a higher accumulation of the inhibitory phosphorylation (Figure S5b).

## Materials and Methods

### Preparation of plasmids

Human CaMKII-α (Uniprot_ID: Q9UQM7) and human CaMKII-β (Uniprot_ID: Q13554) were cloned into the pEGFP-C1 vector backbone (Clontech, Mountain View, CA), after modifying the vector to contain a biotinylation sequence (Avitag, GLNDIFEAQKIEWHE) followed by a linker (GASGASGASGAS) at the N-terminus of mEGFP. CaMKII-α or CaMKII-β was cloned at the C-terminus of mEGFP, with a linker sequence (PreScission protease site: LEVLFQGP) separating the mEGFP tag from the coding sequence of CaMKII-α/CaMKII-β (i.e., the final construct is organized as Avitag-linker-mEGFP-linker-CaMKII-α/CaMKII-β). These constructs were then used as a template to produce the other CaMKII-variants. CaMKII-β* was produced by replacing the linker in CaMKII-α with that from CaMKII-β (Uniprot_ID: Q13554) (Figure S1). mCherry-tagged variants were generated by replacing the mEGFP tag with mCherry in these constructs. pET21a-BirA was a gift from Alice Ting (Addgene plasmid # 20857). BirA was cloned into the pSNAP_f_ vector (New England Biolabs, MA) after modifying the vector backbone to remove the SNAP-tag. All constructs with large domain insertions and deletions were made using standard protocols for Gibson assembly (New England Biolabs, MA). All point mutants used were generated using the standard Quikchange protocols (Agilent Technologies, Santa Clara, CA).

### Tissue culture and DNA transfection

HEK 293T cells (obtained from UC Berkeley cell culture facility) were grown in Dulbecco’s Modified Eagle Medium + GlutaMaX (DMEM, Gibco, Thermo Fisher) that is supplemented with 10% FBS (Avantor Seradigm, VWR, Radnor, Pennsylvania), Antibiotic-Antimycotic (AA, Thermo Fisher) at 100X dilution and 20 mM HEPES buffer and maintained at 37°C under 5% CO_2_. Transient transfection of CaMKII variants were done using the standard calcium phosphate protocol (Wigler et al., 1977). Briefly, CaMKII plasmids (200 ng/400 ng/800 ng depending on the construct) were mixed with 6 μg of empty pcDNA3.1 vector and 1 μg of BirA. This DNA mix was then diluted with ddH_2_O (10X), 250 mM CaCl_2_ was added and the mixture was allowed to sit for 15 min at room temperature. Following this incubation, 2X HBS buffer (50 mM HEPES, 280 mM NaCl, 1.5 mM Na_2_HPO_4_, pH 7.1) was added to it dropwise and mixed thoroughly by reverse pipetting. This mixture was then added to the HEK 293T cells and cells were allowed to express the protein for 18-20 hours before harvesting.

### Preparation of flow cells for single-molecule microscopy

All single-molecule experiments were performed in flow chambers (sticky-Slide VI 0.4, Ibidi, Planegg, Germany) that were assembled with functionalized glass substrates (Ibidi glass coverslips, bottom thickness 170 µm+/–5 µm). The glass substrates were first cleaned using 2% Hellmanex III solution (Hellma Analytics) for 30 min, followed by a 30 minutes sonication in 1:1 mixture (vol/vol) of isopropanol:water. The glass substrates were then dried with nitrogen and cleaned for another 5 minutes in a plasma cleaner (Harrick Plasma PDC-32 G, Ithaca, NY). These cleaned glass substrates were used to assemble the flow chambers immediately after plasma cleaning. After assembly, the glass substrates were treated with a mixture of Poly-L-lysine PEG and PEG-Biotin (1000:1, both at 1 mg/mL) for 30 minutes (SuSoS, Dübendorf, Switzerland). The glass substrates were then washed with 2 mL of Dulbecco’s phosphate-buffered saline (DPBS, Gibco, Thermo Fisher). Streptavidin (Sigma-Aldrich, S0677) was added to these glass substrates at a final concentration of 0.1 mg/mL and incubated for 30 min. Following incubation, excess streptavidin was washed away using 2 mL of DPBS and these assembled flow chambers with streptavidin-coated glass substrates were used for all our single-molecule experiments.

### Cell lysis and pulldown of biotinylated CaMKII in flow chambers

CaMKII variants were allowed to express for 18-20 hours before they were harvested. The co-expression of the *E. coli* biotin ligase, BirA, with the CaMKII variants bearing an Avitag, results in the biotinylation of CaMKII in HEK 293T cells. After harvesting, the cells were lysed in a lysis buffer containing 25 mM Tris at pH 8, 150 mM KCl, 1.5 mM TCEP-HCl, 1% protease inhibitor cocktail (P8340, Sigma), 0.5% phosphatase inhibitor cocktail 2 (P0044, Sigma) and 3 (P5726, Sigma), 50 mM NaF, 15 μg/ml benzamidine, 0.1 mM phenylmethanesulfonyl fluoride and 1% NP-40 (Thermo Fisher). The cell lysate was then diluted 100-200 times in DPBS and 100 μL of this diluted cell lysate was added to a well in the flow chamber for 1 min, before washing it out with 1 mL of DPBS. During this incubation, the biotinylated mEGFP-CaMKII variants were immobilized on the surface of the functionalized glass substrates, owing to the streptavidin-biotin interaction. A buffer exchange was then done in the flow chamber to replace the DPBS with the gel filtration buffer (25 mM Tris, 150 mM KCl, 1.5 mM TCEP, pH 8).

### Activation of CaMKII on glass substrates

The glass-immobilized CaMKII holoenzymes were activated by flowing an activation buffer (with a final concentration of 100 μM CaCl_2_, 10 mM MgCl_2_, 500 μM ATP in the gel filtration buffer and the CaM concentration varies between 0.02 - 5 μM CaM depending on the experiment) into the flow chambers for 60 min. Following this incubation, the activation buffer was washed out using 2 mL of the gel filtration buffer.

### Immunofluorescence assay with phosphospecific antibodies

The autophosphorylation status of the activated CaMKII holoenzyme at the activating (Thr 286) and the inhibitory (Thr 305/306) sites was estimated using phosphospecific antibodies for pThr 286 (Abcam: ab171095) and pThr 305/306 (PhosphoSolutions: p1005-306). The immobilized, activated CaMKII holoenzyme was incubated with a mixture of these two phosphospecific primary antibodies (in a 1:1 ratio using a 1:500 dilution in 5% (w/v) BSA) for 45 min. Subsequently, excess primary antibodies were washed out with 3 mL DPBS. This was followed by a 30-minutes incubation with a 1:1 mix of Alexa-labeled secondary antibodies (1:1000 dilution in 5% BSA), that are complementary to the primary antibodies used. Anti-mouse secondary antibody labeled with Alexa-594 (Cell Signaling Technology) was used for pThr 286-specific primary antibody; anti-rabbit secondary antibody labeled with Alexa-647 (Cell Signaling Technology, Danvers, MA) was used for the pThr 305/306-specific primary antibody. After incubation with the secondary antibody, the flow chambers were washed again using 3 mL of DPBS and the samples were imaged using Total Internal Reflection Fluorescence (TIRF) microscopy.

### Validation of the primary antibodies

The two primary antibodies for pThr 286 and pThr 305/306 do not cross-react with each other’s target sites, as shown by mutations in their respective epitopes (T^286^ is mutated to A^286^ and the epitope for the Thr 305/306 primary antibody is changed from T^305^T^306^MLATRNFS to A^305^V^306^I) in CaMKII-α (Figure S2). These primary antibodies also do not show any reactivity with the unactivated forms of CaMKII-α or CaMKII-β, as pulled down from the cell lysate and before any treatment with activation buffer (data not shown). Additionally, the primary antibody incubation time was optimized so that the phosphosites were nearly saturated by their respective antibodies and we chose an incubation time to be 45 minutes in all our assays (data not shown).

### Phosphatase assay when the kinase is active

To test how the presence of phosphatase affects the phosphorylation status of CaMKII when the kinase is active, increasing amounts of λ-phosphatase (200-800 units, New England Biolabs) and 1 mM MnCl_2_ was added into samples of glass-immobilized CaMKII, in the presence of activation buffer containing 5 μM Ca^2+^/CaM. 1 unit of λ-phosphatase is defined as the amount of enzyme that hydrolyzes 1 nmol of p-nitrophenyl phosphate in 1 minute at 30°C (New England Biolabs). After 45 minutes of incubation, the flow chambers were washed with 2 mL DPBS to remove the λ-phosphatase and the activation buffer. The phosphorylation of Thr 286 and Thr 305/306 was then examined using the immunofluorescence assay described above. No further reduction in phosphorylation was observed upon adding more than 400 units of λ-phosphatase and this is considered as a saturating amount of phosphatase.

### Phosphatase assay when the kinase activity is switched off

To test the sensitivity of the two autophosphorylation sites to phosphatases in the absence of kinase activity, activated CaMKII was treated with two different phosphatases, λ-phosphatase (New England Biolabs) and PP1α (EMD Millipore, Burlington, MA). A phosphatase buffer that contains 1 mM MnCl_2_ and 400-800 units of λ-phosphatase or 800 units of PP1α was added to the flow chambers displaying activated CaMKII, after the activation buffer has been washed out. 1 unit of PP1α is defined as the amount of enzyme that releases 1 nmol phosphate per minute from the phosphorylated substrate DiFMUP (6,8-difluoro-4-methylumbelliferyl phosphate) (EMD Millipore). The dephosphorylation reactions were carried out for defined periods of time (3, 15, or 30 min), following which the phosphatase buffer was washed away with 2 mL DPBS and the autophosphorylation states were examined using the immunofluorescence assay described above.

### Phosphatase assay in solution

The HEK 293T cell lysate was diluted by 1:100-200. CaMKII was activated in this diluted lysate (in solution) by adding an activation buffer containing 5 μM Ca^2+^/CaM and 5 mM Tris-buffered TCEP (Sigma-Aldrich). After activation for 45 min, 100 μM staurosporine (Abcam, ab120056) was added to this reaction mix to inhibit the kinase activity. After inhibiting CaMKII for 10 minutes, saturating amounts of λ-phosphatase (400 units) was added to the solution. Samples were pulled down after 3 or 15 minutes of incubation and the phosphorylation status on Thr 286 and Thr 305/306 was measured. We verified that 100 μM staurosporine was efficient in switching off kinase activity by activating CaMKII-α/β* in the presence of the inhibitor. This treatment completely abolished any autophosphorylation of Thr 286 or Thr 305/306 (Figure S7g).

### Thr 286-phosphorylation recovery assay for CaMKII priming

Activated samples of glass-immobilized CaMKII were treated with saturating amounts of λ-phosphatase and 1 mM MnCl_2_ for 3-5 minutes, followed by washing away of the phosphatase buffer with 2 mL of DPBS. This sample was then incubated with an activation buffer (see details above) containing 25 nM Ca^2+^/CaM for 30 min. The activation buffer was aspirated out with 2 mL DPBS and the phosphorylation status at the activating and inhibitory sites were measured.

### Single-molecule Total Internal Reflection Fluorescence (TIRF) microscopy

Single-particle total internal reflection fluorescence images were acquired on a Nikon Eclipse Ti-inverted microscope equipped with a Nikon 100x 1.49 numerical aperture oil-immersion TIRF objective, a TIRF illuminator, a Perfect Focus system, and a motorized stage. Images were recorded using an Andor iXon electron-multiplying charge-coupled device camera. The sample was illuminated using the LU-N4 laser unit (Nikon, Tokyo, Japan) with solid state lasers for the 488 nm, 561 nm and 640 nm channels. Lasers were controlled using a built-in acousto-optic tunable filter (AOTF). The 405/488/561/638 nm Quad TIRF filter set (Chroma Technology Corp., Rockingham, Vermont) was used along with supplementary emission filters of 525/50m, 600/50m, 700/75m for 488 nm, 561 nm, 640 nm channel, respectively. 3-color image acquisition was performed by computer-controlled change of illumination and filter sets at 42 different positions from an initial reference frame, so as to capture multiple non-overlapping images. Image acquisition was done using the Nikon NIS-Elements software. mEGFP-CAMKII, Alexa-594-labeled anti-Thr 286 antibody, and Alexa-647-labeled anti-Thr 305/306 antibody were imaged by illuminating 488 nm laser set to 5.2 mW, 561 nm laser set to 6.9 mW, and 640 nm laser set to 7.8 mW, respectively. The laser power was measured with the field aperture fully opened. Images were acquired using an exposure time of 80 milliseconds for 488 nm and 561 nm, and an exposure time of 100 milliseconds for 640 nm. The only exception was for acquiring mCherry single-molecule images, where an exposure of 200 milliseconds was used. Epifluorescence images were also acquired using a Nikon Eclipse Ti-inverted microscope with a Nikon 20x objective and an Andor iXon electron-multiplying charge-coupled device camera. A mercury arc lamp (Lumencor Tech., Beaverton, OR) was used for epifluorescence illumination at 488 and 561 nm.

### Analyses of single-molecule TIRF data

Individual single particles in all three channels were detected and localized using the single particle tracking plugin TrackMate in ImageJ (Jaqaman et al., 2008). The particles were localized with the Difference of Gaussian (DoG) detector with an initial diameter set to 6 pixels. The detection threshold value in TrackMate is set to 12 or 15 depending on the wavelength at which the image is acquired. The particles outside the center area of 350 x 350 pixel^2^ were excluded due to heterogeneous TIRF illumination. No further filtering processes was applied in TrackMate.

Intensity distribution and fraction of holoenzymes showing any detectable phosphorylation were analyzed using custom in-house software written in MATLAB. For analyzing the fraction of holoenzymes that show any detectable phosphorylation at a given phosphosite, the XY-coordinates for each particle were extracted from the TrackMate data. For each mEGFP-CAMKII, distances to all the antibody particles were computed, and colocalization was identified if the inter-particle distance was below 2 pixels. The fraction of CAMKII holoenzymes with detectable phosphorylation was computed as a ratio of the number of CAMKII colocalized with antibody to the total number of CAMKII holoenzymes. In a typical case, false-positive colocalization due to coincidentally neighbored particles was less than 0.5%, as determined from analyzing multiple, unrelated sets of images with comparable particle density. The intensity values acquired from TrackMate statistics data for the pThr 286 and pThr 305/306 antibody-channels were used to calculate an intensity histogram, for the colocalized (i.e., corresponding to holoenzymes that show detectable phosphorylation) and the non-colocalized spots (i.e., corresponding to holoenzymes that do not show any detectable phosphorylation with intensity values in the phosphospecific antibody channel being zero). This results in scaling of the area under the histogram for the antibody intensity by a factor that takes into account the fraction of holoenzymes that show no detectable phosphorylation. A bin width of 500 was used for computing the histogram. After calculating the distribution, we plotted the intensity histogram for the colocalized spots only.

For analysis of samples where mEGFP-CaMKII-α and mCherry-CaMKII-β* were co-transfected (i.e., α:β* ratio equals 3:1 and 1:1), we first identified the heterooligomeric holoenzyme population (∼80% of the total holoenzymes, data not shown) and then calculated what fraction of these heterooligomers showed detectable inhibitory phosphorylation. An intensity histogram was also plotted for the pThr 305/306 signal for these holoenzymes, as described above.

## Acknowledgments

We thank members of the Kuriyan lab, especially Laura Nocka for helpful discussions and Miles Tuncel for help with cloning and experimental setup. pET21a-BirA was a gift from Alice Ting (Addgene plasmid # 20857). We thank Darren McAffee for helpful discussions regarding data analysis. MB thanks NIGMS (K99 GM 126145) for funding.

