## Supplemental Figures for "Flexible linkers in CaMKII control the balance between activating and inhibitory autophosphorylation"

### Figure S1

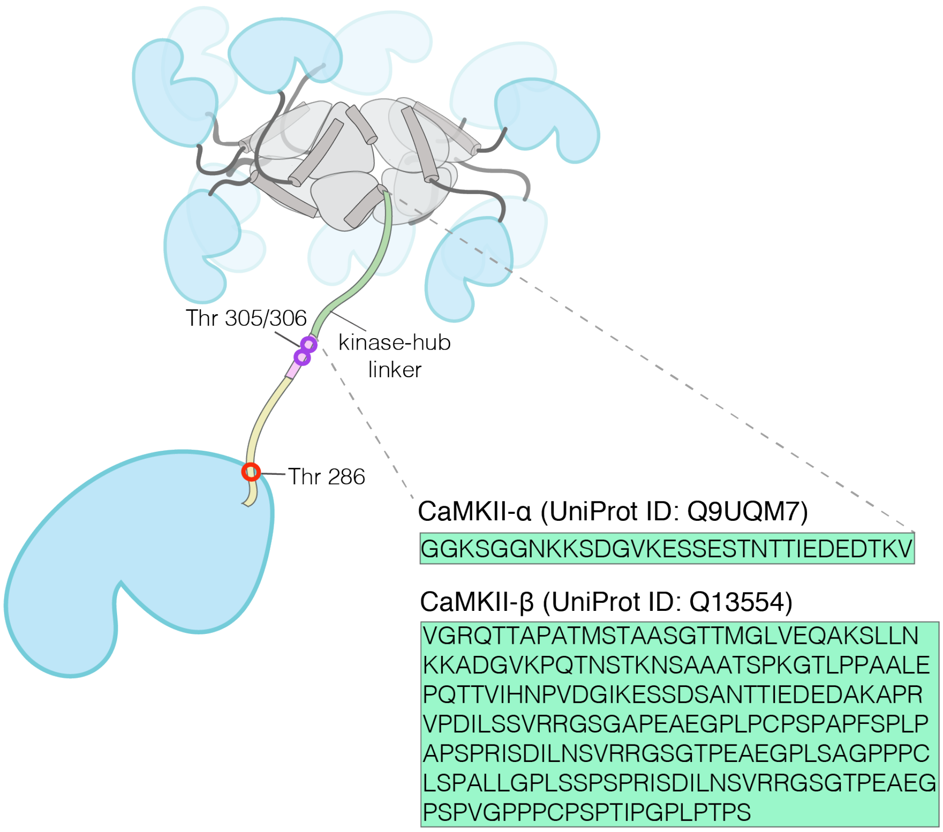


**Figure S1**: Amino acid sequences for the kinase-hub linker in human CaMKII-α and CaMKII-β.

### Figure S2

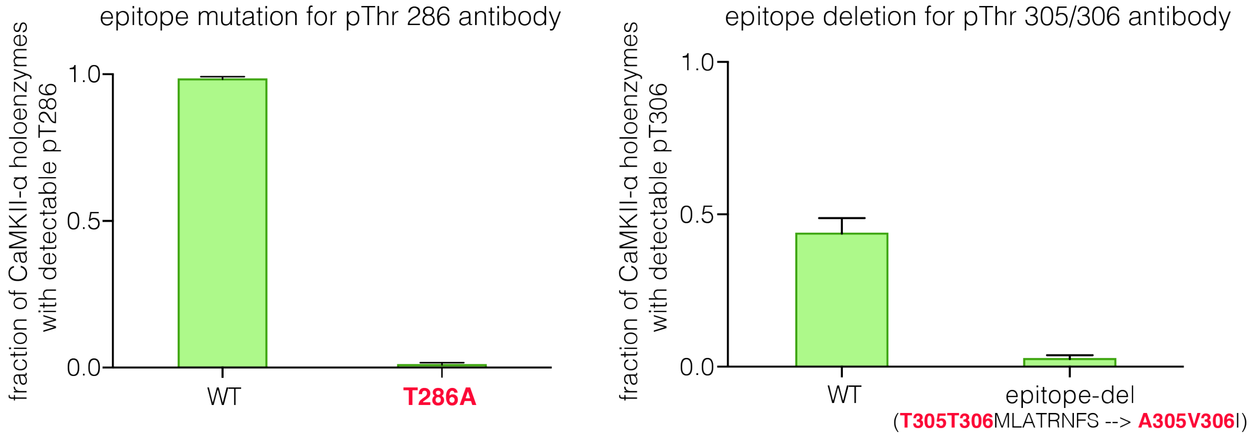


**Figure S2**: Validation of phosphospecific antibodies. Mutation/deletion of epitopes for pThr 286 (left) and pThr 305/306 (right)-specific antibodies leads to no detection of phosphosignal at the respective sites.

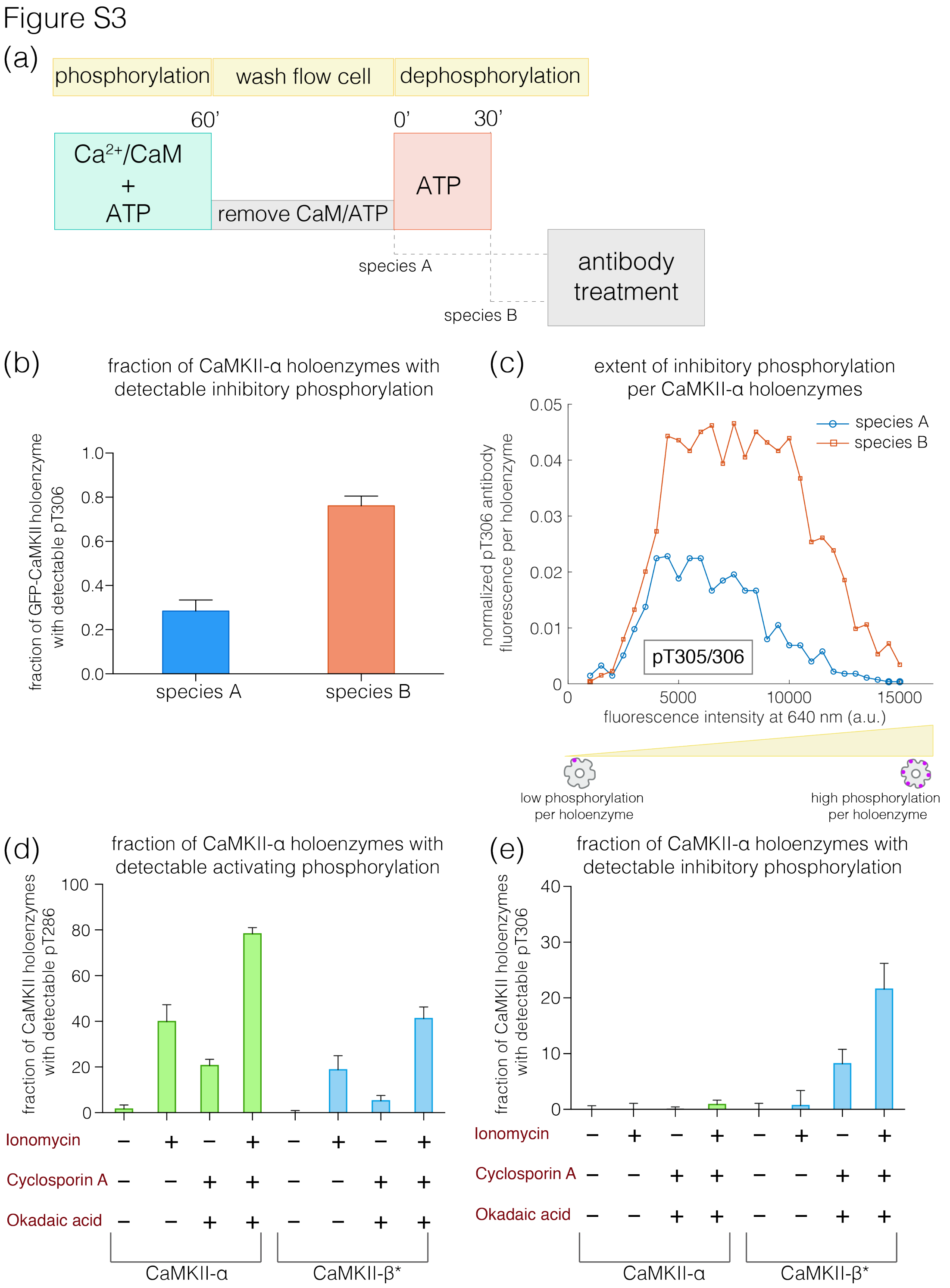


**Figure S3**: (a) Schematic diagram showing the experimental set up. CaMKII was activated, followed by a wash to remove the components of the activation buffer. The pre-activated CaMKII (species A) is then treated with $Mg^{2+}$-ATP for 30 minutes to generate species B. (b) Fraction of CaMKII-α holoenzymes with detectable phosphorylation at Thr 305/306 is plotted for each species. (c) Intensity distribution of pThr 305/306 (640 nm) for species A and species B with detectable phosphorylation. The area under each plot is scaled by the fraction of holoenzymes that show no detectable phosphorylation under that condition (see Methods for details of normalization). (d) Autophosphorylation status of CaMKII after activation in cells. Fraction of CaMKII-α/β* that shows detectable phosphorylation at Thr 286 is plotted for different conditions. (e) Fraction of CaMKII-α/β* that shows detectable phosphorylation at Thr 305/306 is plotted for different conditions. +/- depicts the presence or absence of ionomycin and/or phosphatase inhibitors.

### Figure S5

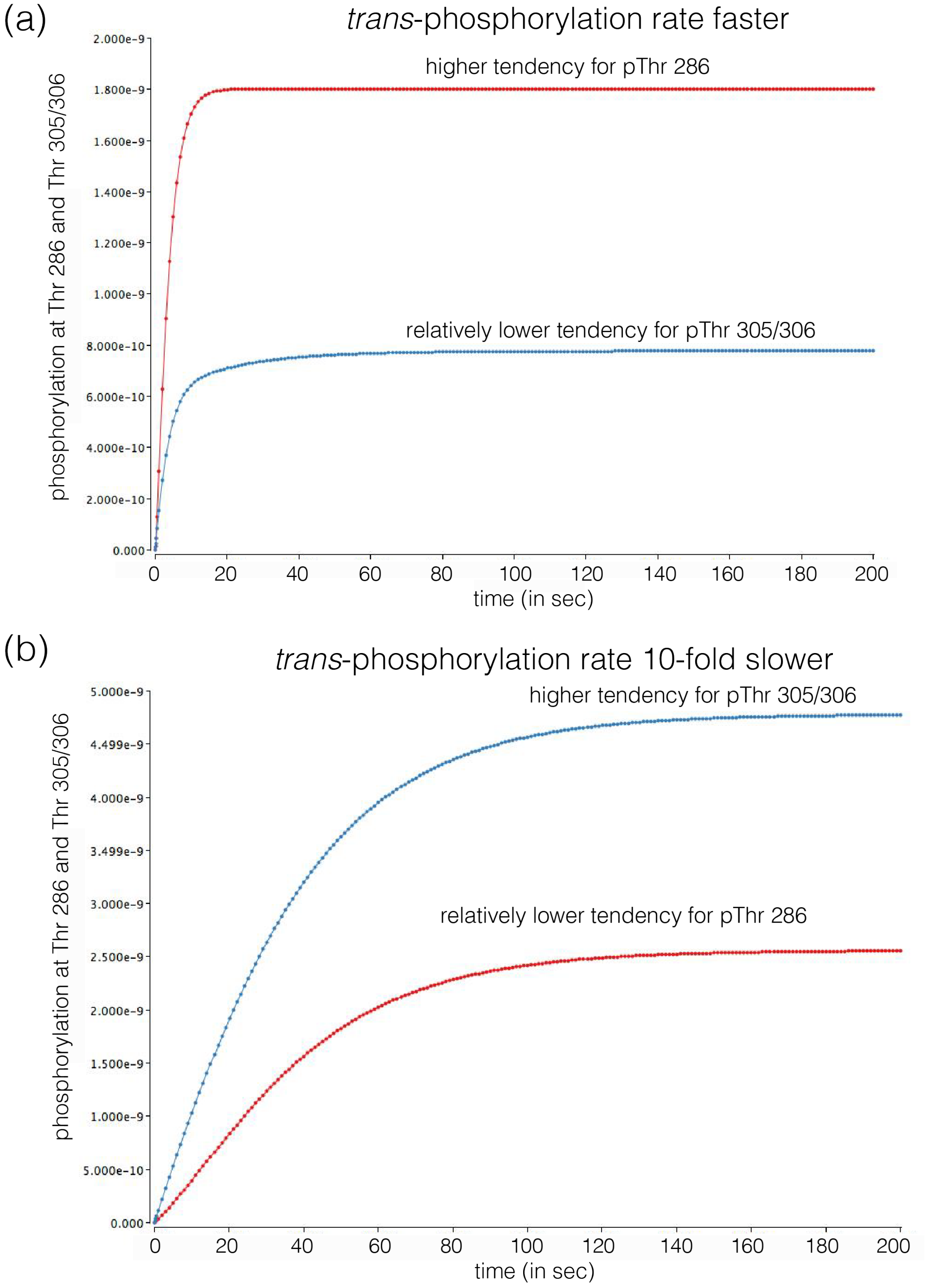


**Figure S5**: Results from simulations of a simple kinetic model for CaMKII using Berkeley Madonna. Plot showing the production of all species bearing pThr 286 or pThr 305/306 over simulation time, when the linker-length is short with faster rates of *trans*-autophosphorylation (a)

and when the linker-length is long and the rates of *trans*-autophosphorylation are 10-fold slower (b).

Figure S7

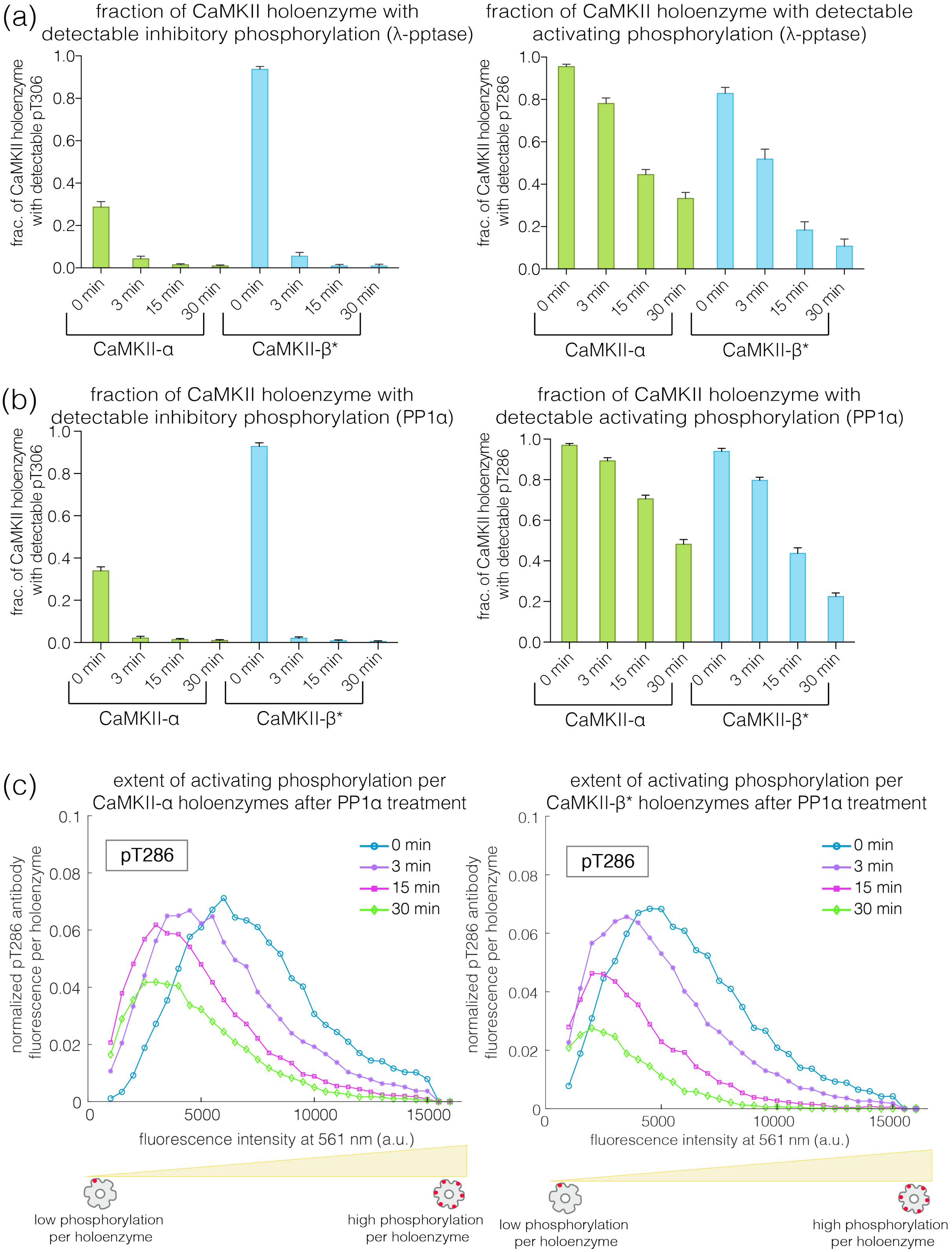
(a)
fraction of CaMKII holoenzyme with detectable inhibitory phosphorylation (λ-pptase)
frac. of CaMKII holoenzyme with detectable pT306
fraction of CaMKII holoenzyme with detectable activating phosphorylation (λ-pptase)
frac. of CaMKII holoenzyme with detectable pT286
0 min
3 min
15 min
30 min
CaMKII-α
CaMKII-β*
(b)
fraction of CaMKII holoenzyme with detectable inhibitory phosphorylation (PP1α)
fraction of CaMKII holoenzyme with detectable activating phosphorylation (PP1α)
(c)
extent of activating phosphorylation per CaMKII-α holoenzymes after PP1α treatment
extent of activating phosphorylation per CaMKII-β* holoenzymes after PP1α treatment
normalized pT286 antibody fluorescence per holoenzyme
pT286
fluorescence intensity at 561 nm (a.u.)
low phosphorylation per holoenzyme
high phosphorylation per holoenzyme

#### Figure S7

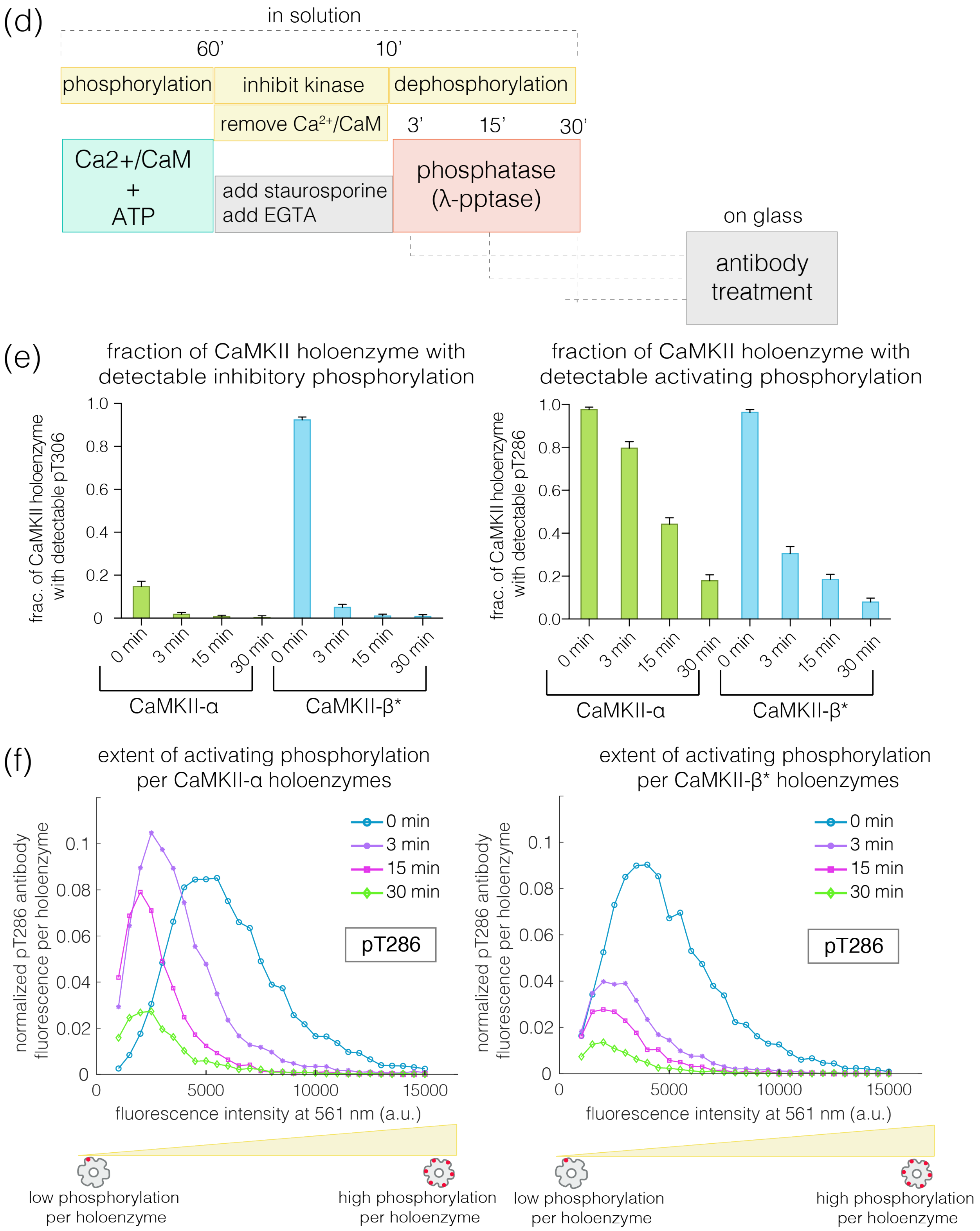
(d)
in solution
60’
10’
phosphorylation
inhibit kinase
dephosphorylation
remove Ca2+/CaM
3’
15’
30’
Ca2+/CaM + ATP
add staurosporine
add EGTA
phosphatase (λ-pptase)
on glass
antibody treatment
(e)
fraction of CaMKII holoenzyme with detectable inhibitory phosphorylation
frac. of CaMKII holoenzyme with detectable pT306
fraction of CaMKII holoenzyme with detectable activating phosphorylation
frac. of CaMKII holoenzyme with detectable pT286
0 min
3 min
15 min
30 min
CaMKII-α
CaMKII-β*
(f)
extent of activating phosphorylation per CaMKII-α holoenzymes
extent of activating phosphorylation per CaMKII-β* holoenzymes
normalized pT286 antibody fluorescence per holoenzyme
fluorescence intensity at 561 nm (a.u.)
pT286
low phosphorylation per holoenzyme
high phosphorylation per holoenzyme

Figure S7

(g)

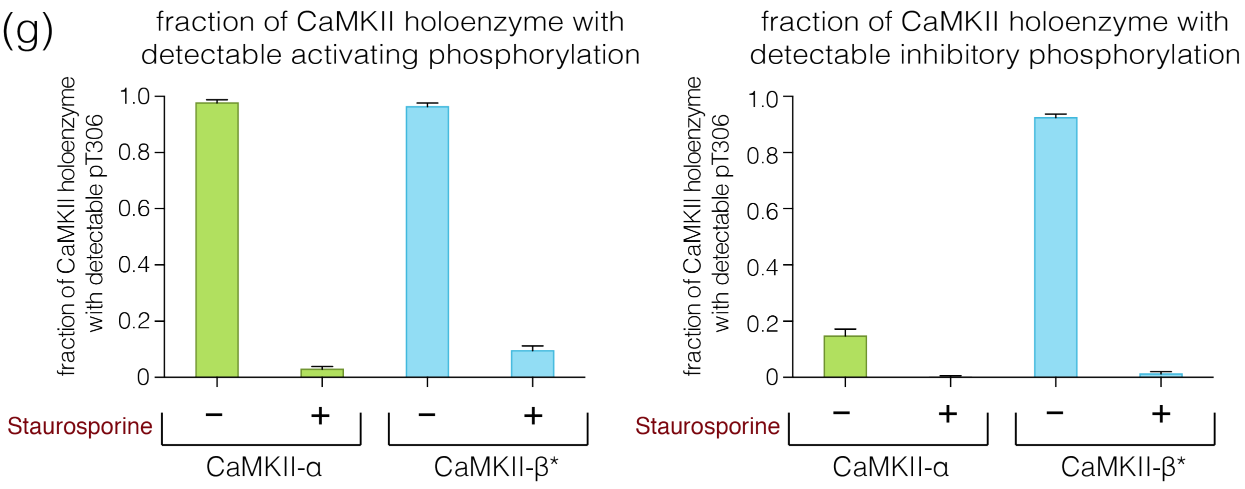


**Figure S7**: Effect of phosphatases on dephosphorylation kinetics when kinase activity is switched off. (a) Fraction of CaMKII-α/β* holoenzymes with detectable phosphorylation at Thr 305/306 (left) and Thr 286 (right) after 0, 3, 15, or 30 minutes of treatment with λ-phosphatase, when the kinase activity is switched off. (b) Same as (a), but PP1α phosphatase was used. (c) Intensity distribution for pThr 286 (561 nm) for CaMKII-α (left panel) and CaMKII-β* (right panel) holoenzymes with detectable phosphorylation at Thr 286 after 0, 3, 15, or 30 minutes of PP1α treatment. The area under each plot is scaled by the fraction of holoenzymes that show no detectable phosphorylation under that condition (see Methods for details of normalization). (d) Schematic diagram depicting the experimental design to measure dephosphorylation kinetics in-solution using λ-phosphatase. (e) Fraction of CaMKII-α/β* holoenzymes with detectable phosphorylation at Thr 305/306 (left) and Thr 286 (right) after 0, 3, 15, or 30 minutes of treatment with λ-phosphatase in-solution, when the kinase activity is switched off by staurosporine. (f) Intensity distribution for pThr 286 (561 nm) for CaMKII-α (left panel) and CaMKII-β* (right panel) holoenzymes with detectable phosphorylation at Thr 286 after 0, 3, 15, or 30 minutes of λ-phosphatase treatment. The area under each plot is scaled by the fraction of holoenzymes that show no detectable phosphorylation under that condition (see Methods for details of normalization). (g) Fraction of CaMKII-α/β* holoenzymes with detectable phosphorylation at Thr 286 (left) and Thr 305/306 (right) after 60 minutes of treatment with the activation buffer (see Methods) in the absence and presence of 100 μM staurosporine.

#### Figure S8

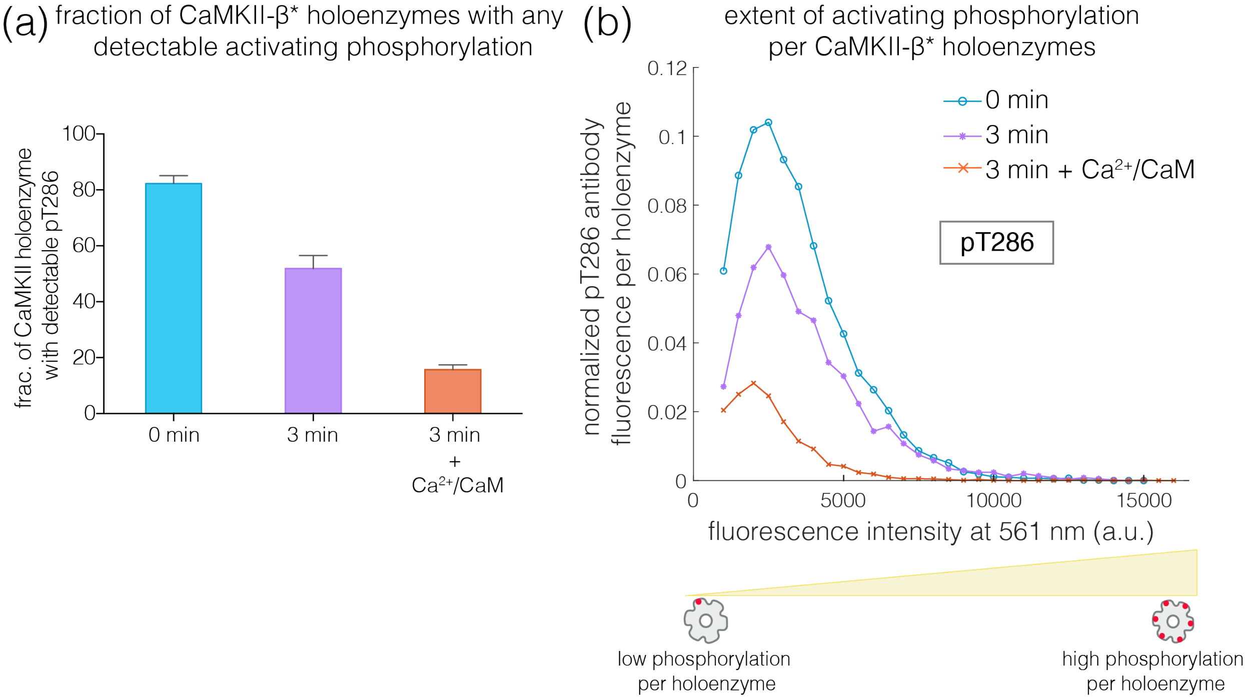


**Figure S8**: Effect of addition of $Ca^{2+}$/CaM on the rates of dephosphorylation at the autonomy site. (a) Fraction of CaMKII-β* holoenzymes with detectable phosphorylation at Thr 286, upon 3 minutes of λ-phosphatase treatment in the absence and presence of 5 μM $Ca^{2+}$/CaM. (b) Intensity distribution of pThr 286 (561 nm) for CaMKII-β* holoenzymes with detectable phosphorylation, under the same conditions as in (a). The area under each plot is scaled by the fraction of holoenzymes that show no detectable phosphorylation under that condition (see Methods for details of normalization).
